# NuMA is a mitotic adaptor protein that activates dynein and connects it to microtubule minus ends

**DOI:** 10.1101/2024.08.30.610456

**Authors:** Sabina Colombo, Christel Michel, Silvia Speroni, Felix Ruhnow, Maria Gili, Claudia Brito, Thomas Surrey

## Abstract

Nuclear mitotic apparatus protein (NuMA) is indispensable for the mitotic functions of the major microtubule minus-end directed motor cytoplasmic dynein 1. NuMA and dynein are both essential for correct spindle pole organization. How these proteins cooperate to gather microtubule minus ends at spindle poles remains unclear. Here we use microscopy-based *in vitro* reconstitutions to demonstrate that NuMA is a dynein adaptor, activating processive dynein motility together with dynein’s cofactors dynactin and Lissencephaly-1 (Lis1). Additionally, we find that NuMA binds and stabilizes microtubule minus ends, allowing dynein/dynactin/NuMA to transport microtubule minus ends as cargo to other minus ends. We further show that the microtubule-nucleating γ-tubulin ring complex (γTuRC) hinders NuMA binding. This shows that either γTuRC needs to be released or microtubules need to be severed to generate free minus ends before they can be incorporated into spindle poles by dynein/dynactin/NuMA. These results provide new mechanistic insight into how dynein, dynactin, NuMA and Lis1 cooperate to organize spindle poles in cells.

## INTRODUCTION

Cytoplasmic dynein 1 (henceforth dynein) is the major microtubule minus-end directed motor protein in animal cells. In interphase, dynein is essential for retrograde transport of a multitude of cargoes, such as vesicles and organelles (Yildiz and Zhao, 2023). During mitosis, it participates in nuclear envelope breakdown and mitotic spindle organization and function (Raaijmakers and Medema, 2014). Human dynein is a large protein complex (≈1.5 MDa) consisting of six distinct polypeptides, each present in duplicate. The C-terminal portion of the heavy chain forms the motor domain consisting of a ring of six AAA domains connected to a microtubule-binding domain by a stalk (Canty et al., 2021). The N-terminal part together with the smaller subunits forms the tail, which serves as a docking site for regulatory components and dynein’s cargo (Reck-Peterson et al., 2018).

When not bound to microtubules, dynein predominantly exists in an auto-inhibited ‘Phi’ conformation (Torisawa et al., 2014; Zhang et al., 2017). The dynein regulator Lis1 can alleviate this auto-inhibition, acting as a molecular wedge that separates the two dynein motor domains (Karasmanis et al., 2023), allowing dynein to bind other interaction partners. Dynein typically interacts with dynactin, another large protein complex (≈1.1 MDa), formed by 23 subunits of 11 different polypeptides, including a central actin-like polymer, the Arp1 filament (Urnavicius et al., 2015). Dynactin serves as a cofactor for virtually all known dynein activities (Canty and Yildiz, 2020). For processive motility, dynein and dynactin need to associate with an adaptor (McKenney et al., 2014; Schlager et al., 2014). Dynein adaptors are coiled coil proteins whose N-terminal part is sandwiched in between dynein and dynactin, thereby stabilizing dynein’s and dynactin’s active conformation (Chowdhury et al., 2015; Chaaban and Carter, 2022). The C-terminal part of adaptors contains the cargo-binding domain (Carter, Diamant and Urnavicius, 2016; Olenick and Holzbaur, 2019). Therefore, in addition to promoting dynein activation, adaptors bind specific cargoes, providing the dynein/dynactin complex with functional versatility (Olenick and Holzbaur, 2019; Canty and Yildiz, 2020).

During cell division, dynein is indispensable for the correct functioning of meiotic and mitotic spindles. One of its important roles is spindle pole focusing, thought to be achieved by motor-driven gathering of microtubule minus ends (Verde et al., 1991; Heald et al., 1996; Sikirzhytski et al., 2014; Hueschen et al., 2019; So et al., 2022). How dynein crosslinks microtubules and transports minus ends towards other minus ends remains however unclear.

Dynein’s pole focusing activity requires its ubiquitous partners dynactin and Lis1 (Wang et al., 2013; Monda and Cheeseman, 2018; So et al., 2022). Moreover, it requires also a mitosis-specific interaction partner, the nuclear mitotic apparatus protein (NuMA). In animal cells, NuMA localizes to the nucleus during interphase. In mitosis, it accumulates at spindle poles to contribute to proper pole organization (Lydersen and Pettijohn, 1980; Maekawa, Leslie and Kuriyama, 1991; Gaglio, Saredi and Compton, 1995; Merdes et al., 1996, 2000; Heald et al., 1997; Hueschen et al., 2017), and it recruits dynein to the cell cortex to ensure correct spindle positioning(Okumura et al., 2018).

NuMA is a homodimeric protein with a long central coiled coil (≈210 nm) (Harborth, Weber and Osborn, 1995; Harborth et al., 1999). Its N-terminal Hook domain binds the dynein light intermediate chain, and is adjacent to a CC1-box-like motif, conserved among various dynein adaptors (Renna et al., 2020). A Spindly-like motif has also been identified (Okumura et al., 2018; Tsuchiya et al., 2020), which may promote the association with dynactin’s pointed end (Gama et al., 2017; Lee et al., 2020). NuMA co-immunoprecipitates with dynein and dynactin in *Xenopus* egg extract (Merdes et al., 1996), and its first 505 amino acids are sufficient for cortical recruitment of dynein in human cells (Okumura et al., 2018). Although NuMA has therefore been proposed to function as an activating dynein adaptor (Hueschen et al., 2017; Reck-Peterson et al., 2018; Renna et al., 2020), this has not been directly demonstrated yet.

NuMA’s C-terminal part has been shown to be required for correct spindle organization in human cells (Hueschen et al., 2017; Okumura et al., 2018; Pirovano et al., 2019). It interacts with microtubules through two proposed microtubule binding domains (MTBDs) (Du et al., 2002; Gallini et al., 2016; Chang et al., 2017). Moreover, *in vitro* experiments with purified proteins revealed that the C-terminal part can also support NuMA’s self-assembly into oligomers (Harborth et al., 1999), and trigger phase separation (Sun et al., 2021), which may explain NuMA’s clustering behavior observed in cells (Okumura et al., 2018), and may contribute to passive microtubule crosslinking (Merdes et al., 1996; Nachury et al., 2001; Haren and Merdes, 2002). In human cells, NuMA has also been shown to localize to the minus ends of laser-ablated kinetochore fibers independently of dynein (Hueschen et al., 2017), raising the possibility that NuMA has the intrinsic property of recognizing microtubule minus ends, which has however not been tested directly. The molecular mechanism by which NuMA contributes to dynein’s microtubule minus-end gathering activity remains therefore unclear.

Here, we investigate the interplay between NuMA, dynein and microtubules using total internal reflection fluorescence (TIRF) microscopy-based *in vitro* reconstitution assays with purified proteins. We find that the N-terminal part of NuMA can activate processive dynein motility and that this activation does not only require dynactin but also Lis1. We demonstrate that NuMA’s C-terminal part directly binds microtubules with preference for free minus ends, capping and stabilizing them. Finally, we show that the dynein/dynactin/NuMA complex can transport minus ends of cargo microtubules towards minus ends of other microtubules. These results establish NuMA as an activating dynein adaptor, whose cargo is a microtubule minus end. These results provide mechanistic insight into the molecular mechanism by which dynein, dynactin, NuMA, and Lis1 cooperate to focus spindle poles during mitosis.

## RESULTS

### NuMA is a dynein adaptor that requires Lis1 and dynactin to activate dynein motility

We purified a recombinant N-terminal fragment of human NuMA consisting of its first 705 amino acids (aa), fused to a SNAP-tag that was either labelled with Alexa Fluor 546 or 647 (AF546 or AF647-NuMA^N-term^, Fig. 1 A, Fig. S1 A). This NuMA construct was previously shown to bind to dynein in cells (Kotak, Busso and Gönczy, 2012) and to the dynein light intermediate chain *in vitro* (Renna et al., 2020). Due to the presence of part of NuMA’s predicted coiled coil, NuMA^N-term^ was dimeric as demonstrated by mass photometry (Fig. S1 B). We also purified a recombinant human dynein complex with monomeric EGFP (mEGFP) fused to its heavy chain, and porcine brain dynactin (Fig. S1 A), as described previously (Jha et al., 2017).

**Fig. 1.**
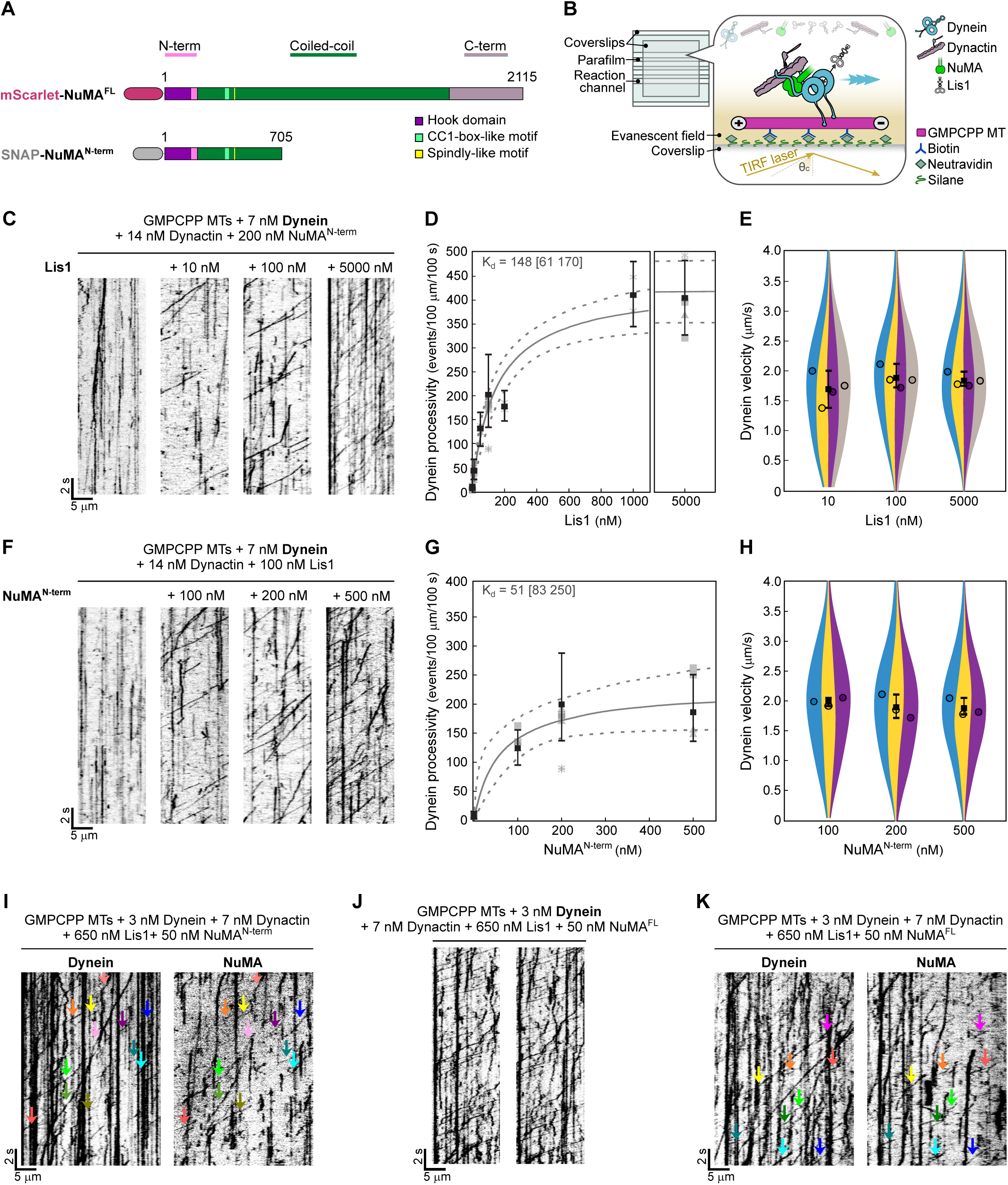
NuMA is a dynein adaptor that requires dynactin and Lis1 to activate dynein motility. **(A)** Schematic of mScarlet-tagged NuMA^FL^ and SNAP-tagged NuMA^N-term^ constructs, indicating the main structural parts of NuMA (N-terminal region, predicted coiled coil, C-terminal tail) and the domains or motifs implicated in dynein-binding (Okumura et al., 2018; Renna et al., 2020). **(B)** Schematic of microscopy flow chambers (left), with functionalized glass surface and components of dynein motility assays (right). **(C)** Representative TIRF microscopy kymographs showing the motility of mEGFP-dynein in the presence of dynactin and AF546-NuMA^N-term^ at different mCherry-Lis1 concentrations. Concentrations as indicated. **(D)** Processive run frequency of mEGFP-dynein (black squares and whiskers: mean ± 95% CI). Each distinct grey symbol (circle, asterisk, square, triangle) refers to an independent replicate; each data point represents the number of events in one field of view of one replicate; *n* = 99, 101, 27, 100, 29, 26, 103 microtubules. Grey curve represents a hyperbolic fit; apparent K_d_ is indicated (dashed lines: 95% confidence interval). **(E)** Velocity distribution of processive mEGFP-dynein runs (mean of medians ± 95% CI, four independent experiments). Each circle represents the median velocity of one replicate; *n* = 760, 3751, 7088 velocities. Protein combination and concentrations in D, E as in C and as indicated. **(F)** Representative kymographs showing the motility of mEGFP-dynein in the presence of dynactin and mCherry-Lis1 at different AF546-NuMA^N-term^ concentrations. Concentrations as indicated. **(G)** Processive run frequency of mEGFP-dynein (mean ± 95% CI). Symbols and curves as in D; *n* = 103, 70, 100, 70 microtubules. **(H)** Velocity distribution of processive mEGFP-dynein runs (mean of medians ± 95% CI, three independent experiments). Each circle represents the median velocity of one replicate; *n* = 1585, 3482, 2649 velocities. Protein combinations and concentrations in G, H as in F, and as indicated. Representative kymographs showing the motility of: **(I)** mEGFP-dynein (left) and AF647-NuMA^N-term^ (right) in the presence of dynactin and Lis1, **(J)** mEGFP-dynein in the presence of dynactin, Lis1, and mScarlet-NuMA^FL^, **(K)** mEGFP-dynein (left) and mScarlet-NuMA^FL^ (right) in the presence of dynactin and Lis1. Arrowheads of same color indicate co-localization in the same processive events. All data refer to motility on biotinylated Atto647N-GMPCPP-microtubules (MTs). Experiments shown in C‒H and J were carried out at 30 °C, and those shown in I and K at 18 °C.

To test whether NuMA can act as a dynein adaptor, we immobilized stabilized Atto647N-labelled microtubules on a glass surface, added AF546-NuMA^N-term^, mEGFP-dynein and dynactin, and observed mEGFP-dynein by TIRF microscopy (Fig. 1 B). Under these conditions, we hardly ever observed processive motility events along microtubules (Fig. 1 C, left kymograph), in contrast to the typical behavior of dynein in the presence of dynactin and an adaptor (McKenney et al., 2014; Schlager et al., 2014). We found that addition of purified human Lis1, that is known to relieve dynein’s auto-inhibition (Qiu, Zhang and Xiang, 2019; Elshenawy et al., 2020; Htet et al., 2020; Marzo, Griswold and Markus, 2020; Karasmanis et al., 2023), was required to trigger dynein to move processively in the presence of NuMA^N-term^ and dynactin. Lis1 increased the number of processive motility events in a dose-dependent manner (Fig. 1 C and D), similarly to what can be observed with other adaptors, such as bicaudal D-related protein 1 (BicDR1) (Zhao, Oten and Yildiz, 2023) or protein bicaudal D homolog 2 (BicD2)N^1‒400^ (Fig. S2 A), which however do not strictly require Lis1(McKenney et al., 2014; Schlager et al., 2014; Schroeder and Vale, 2016; Redwine et al., 2017; Urnavicius et al., 2018; Canty et al., 2023). Increasing Lis1 concentration did not affect dynein velocity (Fig. 1 E), as previously also shown for BicD2 (Jha et al., 2017).

In the absence of NuMA^N-term^, as expected Lis1 did not stimulate processive dynein motility, because it is not an activating adaptor (Fig. 1 F, left kymograph). Increasing the concentration of NuMA^N-term^, while keeping the Lis1 concentration constant, increased the number of processive dynein motility events (Fig. 1 F and G) without affecting dynein velocity (Fig. 1 H). The average dynein velocity, measured at 30 °C across all displayed conditions (Fig. 1 E and H), was ≈1.9 µm s^-1^. This is higher than reported *in vitro* velocities of mammalian dynein (Elshenawy et al., 2020; Htet et al., 2020; Canty et al., 2023; Zhao, Oten and Yildiz, 2023) due to the higher temperature in our experiments (Ruhnow, Zwicker and Diez, 2011; Hong et al., 2016) (Fig. S2 B). We observed no difference between dynein velocities in the presence of NuMA^N-term^ or BicD2N^1‒400^, under the same conditions (Fig. S2 B). Using a relatively low NuMA^N-term^ concentration to reduce fluorescence background, and a relatively high Lis1 concentration, allowed the visualization of AF647-NuMA^N-term^ transport by mEGFP-dynein, in agreement with NuMA’s activating dynein adaptor function (Fig. 1 I, arrowheads).

Next, we purified recombinant full-length human NuMA fused to the fluorescent protein mScarlet (Scarlet-NuMA^FL^) and tested its ability to stimulate processive dynein motility. We found that also NuMA^FL^ was able to activate dynein in the presence of both dynactin and Lis1 (Fig. 1 J), similarly to what was observed with NuMA^N-term^. This indicates that full-length NuMA under our conditions is not, or at least not completely, auto-inhibited, similarly to BicDR1 and Hook3 (Urnavicius et al., 2018; Kendrick et al., 2019), and differently from the adaptors BicD1/2, Spindly, and JNK-interacting protein 3 (JIP3) (Liu et al., 2013; Carter, Diamant and Urnavicius, 2016; D’amico et al., 2022; Singh et al., 2023). Lack of clear auto-inhibition was further supported by the elongated appearance of NuMA in negative stain electron microscopy, displaying a length roughly corresponding to the expected contour length of its predicted coiled coil (S1 C) (Harborth, Weber and Osborn, 1995). Additionally, dynein velocities in the presence of NuMA^FL^ or NuMA^N-term^ were similar (Fig. S2 B). mScarlet-NuMA^FL^ could also be observed to be transported by mEGFP-dynein, again in agreement with NuMA’s activating adaptor function (Fig. 1 K, arrowheads).

These results establish NuMA as a new dynein adaptor whose dynein processivity-stimulating activity depends more strongly on the additional presence of Lis1 than that of other dynein adaptors.

### NuMA’s main microtubule binding region is located close to its C-terminus

Next, we purified three recombinant C-terminal fragments of human NuMA fused to mScarlet (Fig. 2 A, Fig. S3 A and B): (i) A long C-terminal fragment comprising aa 1560‒2115 (NuMA^C-term^ ^L^), which contains part of the predicted coiled coil and the entire C-terminal "tail" region previously reported to contain two microtubule binding domains (Du et al., 2002; Gallini et al., 2016; Chang et al., 2017) and a clustering domain (Okumura et al., 2018). (ii) A shorter fragment comprising aa 1882‒2105 (NuMA^C-term^ ^S2^), which contains only part of the "tail", including the reported microtubule binding domains, but lacks the clustering domain (similar or identical to what was previously called NuMA-tail II (Nachury et al., 2001; Wiese et al., 2001; Haren and Merdes, 2002; Forth et al., 2014; Chang et al., 2017) or NuMA C-tail2(Hueschen et al., 2017)). (iii) Another short fragment comprising aa 1701‒1981 (NuMA^C-term^ ^S1^), which contains also part of the "tail", but lacks the most C-terminal reported microtubule binding domain (previously named C-tail 1+2A (Hueschen et al., 2017)).

**Fig. 2.**
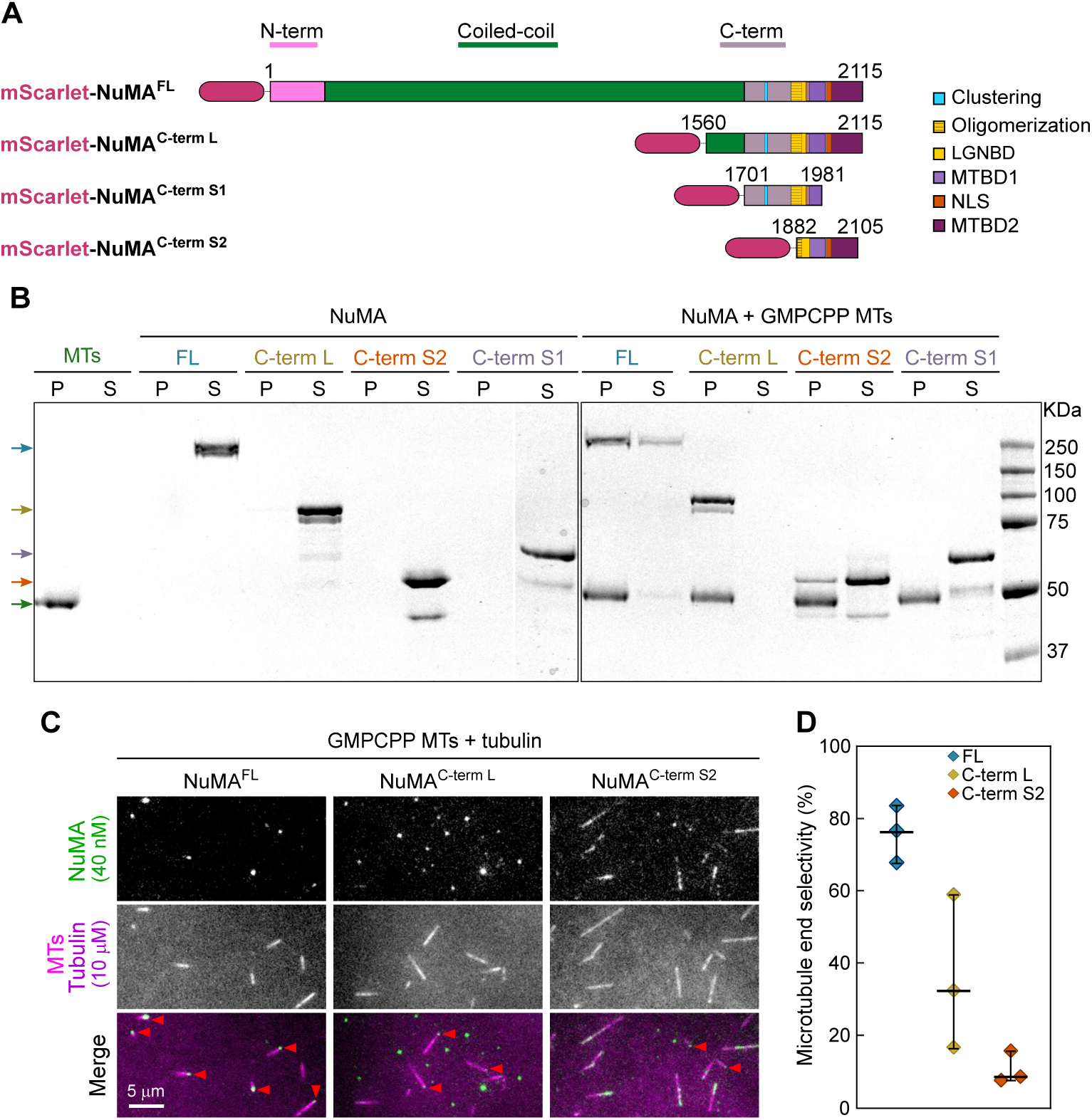
NuMA’s microtubule binding region is located close to its C-terminus. **(A)** Schematic of mScarlet-tagged NuMA^FL^ and C-terminal fragments, indicating the main structural parts of NuMA and functional domains identified in the C-terminal region (microtubule binding domain [MTBD] 1 (Du et al., 2002) and 2 (Gallini et al., 2016; Chang et al., 2017), clustering region (Okumura et al., 2018), oligomerization region (Pirovano et al., 2019), LGN binding domain [LGNBD] (Pirovano et al., 2019), nuclear localization signal [NLS] (Chang et al., 2017)). **(B)** Coomassie Blue-stained SDS-PAGE showing the pellet (P) and supernatant (S) fractions of microtubule co-sedimentation assays. 1 µM mScarlet-tagged NuMA^FL^ or C-terminal fragments were incubated alone or with Atto647N-labelled GMPCPP-microtubules (0.5 µM polymerized tubulin) at 30 °C for 15 min. Upon centrifugation, in the absence of microtubules, all proteins remained in the supernatant (left gel). When incubated with microtubules, they co-sedimented with the microtubule pellet (green arrow) at different degrees. NuMA^C-term^ ^L^ was found exclusively in the pellet; NuMA^FL^ predominantly in the pellet; NuMA^C-term^ ^S2^ partially in the pellet; NuMA^C-term^ ^S1^ entirely in the supernatant. Arrows indicate the expected molecular weight for each NuMA construct. **(C)** Representative TIRF microscopy images of 40 nM mScarlet-tagged NuMA constructs binding to biotinylated Atto647N-labelled GMPCPP-microtubules in the presence of 10 µM Atto647N-tubulin. Arrowheads indicate selective end binding. **(D)** Microtubule end selectivity: percentage of microtubules showing mScarlet signal exclusively at one end, of all microtubules with a mScarlet signal, for mScarlet-NuMA constructs binding to microtubules in the presence of tubulin as shown in C (mean ± SD). Each diamond represents one replicate; *n* = 126, 71, 161 microtubules. All data refer to three independent experiments.

Using mass photometry, we observed that the longer NuMA^C-term^ ^L^ fragment was a dimer, however partly dissociated into monomers, whereas both short fragments lacking any predicted coiled coil were monomers (Fig. S3 C). We could not analyze the oligomerization state of NuMA^FL^, given its low concentration and the presence of detergent in its buffer.

Using a microtubule co-sedimentation assay, we observed that NuMA^FL^, NuMA^C-term^ ^L^ and NuMA^C-term^ ^S2^ bound to GMPCPP-microtubules (Fig. 2 B). The binding of NuMA^C-term^ ^L^ was strongest, followed by NuMA^FL^ and NuMA^C-term^ ^S2^. NuMA^C-term^ ^S2^ binds weaklier to microtubules than the longer constructs, probably because it is monomeric. NuMA^C-term^ ^S1^ did not bind to microtubules under these conditions, in agreement with a previous *in vitro* study suggesting that the major microtubule binding region in NuMA is at its very C-terminus (Chang et al., 2017).

Next, we were interested in observing by TIRF microscopy how the different NuMA constructs bind to microtubules growing from surface-immobilized stabilized GMPCPP-microtubule "seeds" in the presence of tubulin. Remarkably, at the considerably lower NuMA concentrations used here compared to the co-sedimentation assay, NuMA^FL^ and to a lesser extent NuMA^C-term^ ^L^ appeared to bind preferentially to one of the two microtubule ends (Fig. 2 C, arrowheads) which was confirmed by quantifying the selectivity for binding to single ends (Fig. 2 D). NuMA^C-term^ ^L^ and to a lesser extent NuMA^FL^ were also observed to non-specifically adsorb to the surface as what appeared to be clusters of varying size, probably due to the presence of a clustering domain within these constructs (Okumura et al., 2018).

### NuMA’s microtubule binding region preferentially binds to microtubule minus ends and prevents their growth

To determine to which microtubule end NuMA binds with preference, we imaged NuMA^FL^ and the C-terminal NuMA fragments over time as microtubules elongated from the GMPCPP-"seeds" in the presence of free tubulin (Fig. 3 A). In the absence of NuMA, plus and minus ends can easily be distinguished by their different growth speeds (Fig. 3 B). Adding increasing concentrations of NuMA^FL^ slowed down the growth of the slower growing minus ends in a dose-dependent manner (Fig. 3 C and D). In contrast, growth of the faster growing plus ends was largely unaffected in the studied concentration range (which was limited by the solubility of NuMA^FL^). At the highest tested concentration, minus-end growth was completely prevented, with NuMA^FL^ accumulating selectively to these ends, demonstrating that NuMA has a microtubule minus-end binding preference. Only a minor decrease in velocity was observed for the growth of plus ends at the highest NuMA^FL^ concentration.

**Fig. 3.**
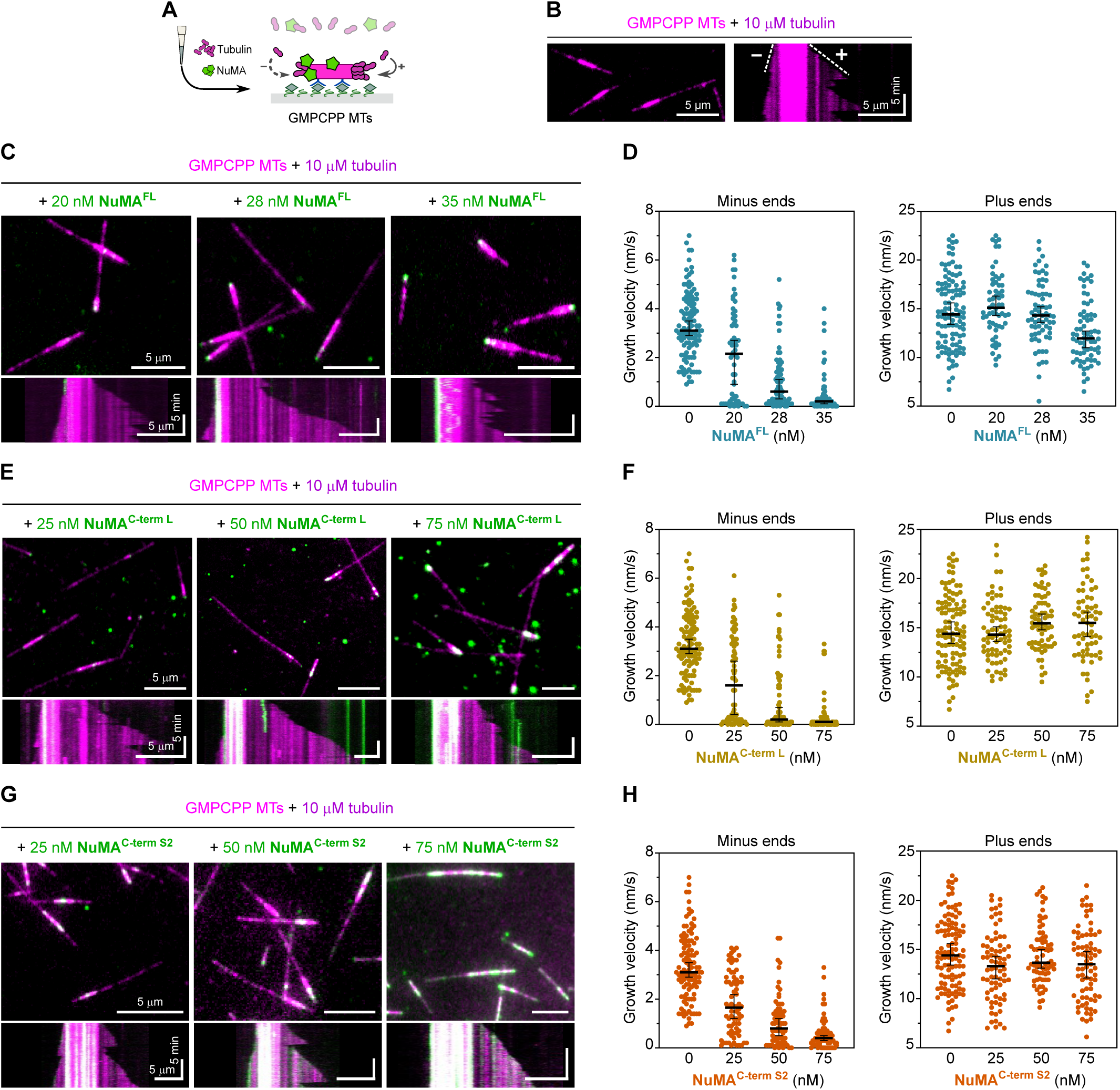
NuMA’s microtubule binding region preferentially binds to microtubule minus ends and prevents their growth. **(A)** Schematic of a TIRF microscopy assay with fluorescent NuMA constructs and tubulin being flowed simultaneously into a channel containing immobilized GMPCPP-microtubule seeds. **(B)** Representative TIRF microscopy image (left) and kymograph (right) showing minus and plus ends (dim magenta) dynamically elongating from a biotinylated Atto647N-labelled GMPCPP-seed (bright magenta) in the presence of Atto647N-tubulin. **(C, E, G)** Representative TIRF microscopy images (top) and kymographs (bottom) showing the growth behavior of microtubule ends (dim magenta) elongating from biotinylated Atto647N-labelled GMPCPP-seeds (bright magenta) in the presence of Atto647N-tubulin and various concentrations of different mScarlet-tagged NuMA constructs (green). **(D, F, H)** Growth velocity distributions of microtubule minus and plus ends related to the experiments shown in C, E, G (median ± 95% CI). Each circle represents the growth velocity of one microtubule end segment; D: *n* = 109, 60, 69, 71 (left plot) and 109, 59, 68, 70 (right plot); p-values: 0.00 for all conditions (left plot) and 0.15, 0.99, 0.00 (right plot). F: *n* = 109, 72, 66, 67 (left plot) and 109, 72, 68, 65 (right plot); p-values: 0.00 for all conditions (left plot) and 1.00, 0.16, 0.55 (right plot). H: *n* = 109, 74, 70, 66 (left plot) and 109, 74, 70, 75 (right plot); p-values: 0.00 for all conditions (left plot) and 0.04, 1.00, 0.11 (right plot). Data are from three independent experiments per condition; p-values were calculated from a Kruskal-Wallis test comparing each condition to the control at 0 nM NuMA.

A similar behavior was observed for NuMA^C-term^ ^L^ (Fig. 3 E and F) and NuMA^C-term^ ^S2^ (Fig. 2 G and H), although the inhibitory effect of the shorter construct NuMA^C-term^ ^S2^ on minus end growth was milder, possibly due to this construct being monomeric instead of dimeric. In contrast, NuMA^C-term^ ^S1^ did not exert a clear effect on microtubule dynamics, not even at elevated concentrations, when it began to weakly bind to GMPCPP-seeds (Fig. S4 A and B).

Taken together, these results show that NuMA’s main microtubule binding domain, located at its C-terminus, is required for selective minus end growth inhibition, and that NuMA^C-term^ ^S2^ is sufficient to exert this effect, even though longer NuMA constructs act more strongly.

### NuMA caps and stabilizes dynamic microtubule minus ends

To exclude that minus-end recognition by NuMA is a GMPCPP-microtubule-specific effect, we performed microtubule pre-elongation experiments (Fig. 4 A and B). We first allowed dynamic microtubules to grow from immobilized GMPCPP-seeds in the presence of tubulin for ≈10 min, before adding NuMA while keeping the tubulin concentration constant. Also here, NuMA^FL^ bound preferentially to microtubule minus ends, stopping their growth. Moreover, it stabilized these minus ends, preventing catastrophe after growth stopped, making them static. In contrast, plus-end growth was again mostly unaffected (Fig. 4 C and D, Video 1). NuMA^C-term^ ^L^ and NuMA^C-term^ ^S2^ exhibited similar effects, with NuMA^C-term^ ^S2^ requiring slightly higher concentrations (Fig. S5). In agreement with our previous observations, NuMA^C-term^ ^S1^ did not display any binding to dynamic microtubule ends (Fig. S4 C).

**Fig. 4.**
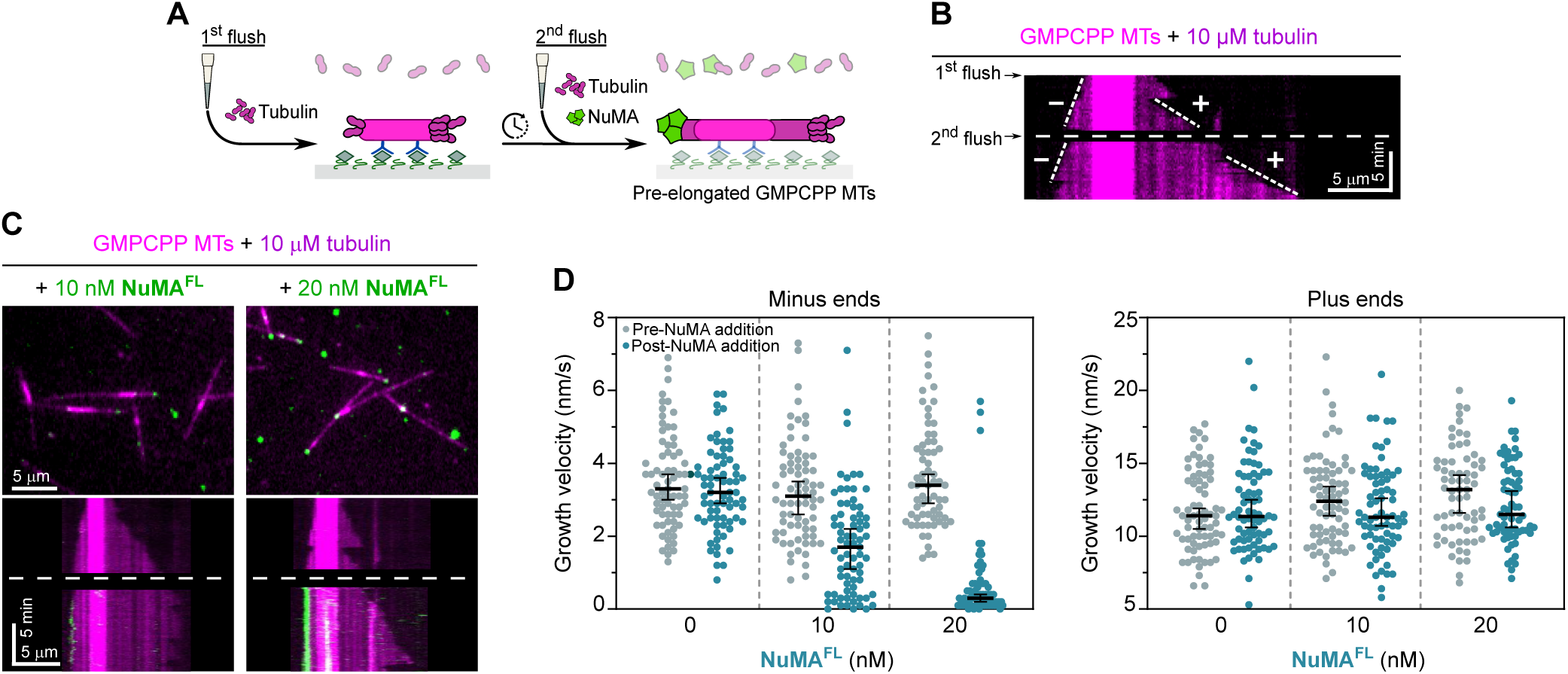
NuMA caps and stabilizes dynamic microtubule minus ends. **(A)** Schematic of a two-flush TIRF microscopy assay: fluorescent tubulin is flowed into a channel containing immobilized GMPCPP-seeds (1^st^ flush); microtubules are allowed to elongate from GMPCPP-seeds for ≈10 min; tubulin is re-added together with fluorescent NuMA constructs (2^nd^ flush). **(B)** Representative kymograph showing a control experiment with microtubule minus and plus ends (dim magenta) dynamically elongating from a biotinylated Atto647N-labelled GMPCPP-seed (bright magenta) in the presence of 10 µM Atto647N-tubulin; the dashed line marks the second flush of tubulin. **(C)** Representative TIRF microscopy images (top) and kymographs (bottom) showing the growth behavior of microtubule ends (dim magenta) elongating from biotinylated Atto647N-labelled GMPCPP-seeds (bright magenta) in the presence of Atto647N-tubulin, before (above the dashed line) or after (below the dashed line) the addition of mScarlet-NuMA^FL^ (green) at different concentrations. Related to Video 1. **(D)** Growth velocity distributions of minus and plus ends elongating from GMPCPP-seeds in the presence of 10 µM Atto647N-tubulin before and after the addition of mScarlet-NuMA^FL^ at different concentrations (median ± 95% CI, three independent experiments). Each circle represents the growth velocity of one microtubule end segment; *n* = 72, 73, 67; p-values: 1.00, 0.00, 0.00 (left plot) and 1.00, 0.59, 0.58 (right plot); p-values were calculated from a Kruskal-Wallis test comparing “pre-NuMA addition” and “post-NuMA addition” velocities for each condition.

In conclusion, NuMA preferentially binds to dynamic microtubule minus ends, stabilizing and capping them at nanomolar NuMA concentrations. These results establish NuMA as a new autonomous minus end capper.

### NuMA does not bind to γTuRC-capped microtubule minus ends

Most microtubules in eukaryotic cells are nucleated by the γ-tubulin ring complex (γTuRC), which naturally caps their minus ends from the start of the growth (Zheng et al., 1995; Moritz et al., 2000; Consolati et al., 2020; Wieczorek et al., 2021; Rai et al., 2024). We therefore tested whether NuMA could bind to the minus ends of γTuRC-nucleated microtubules. We immobilized purified mBFP-labelled and biotinylated human γTuRC on a functionalized glass surface and observed by TIRF microscopy how microtubules were nucleated and grew in the presence of NuMA^FL^ (Fig. 5 A). Most microtubules were nucleated by γTuRC, but some were also nucleated spontaneously in solution.

**Fig. 5.**
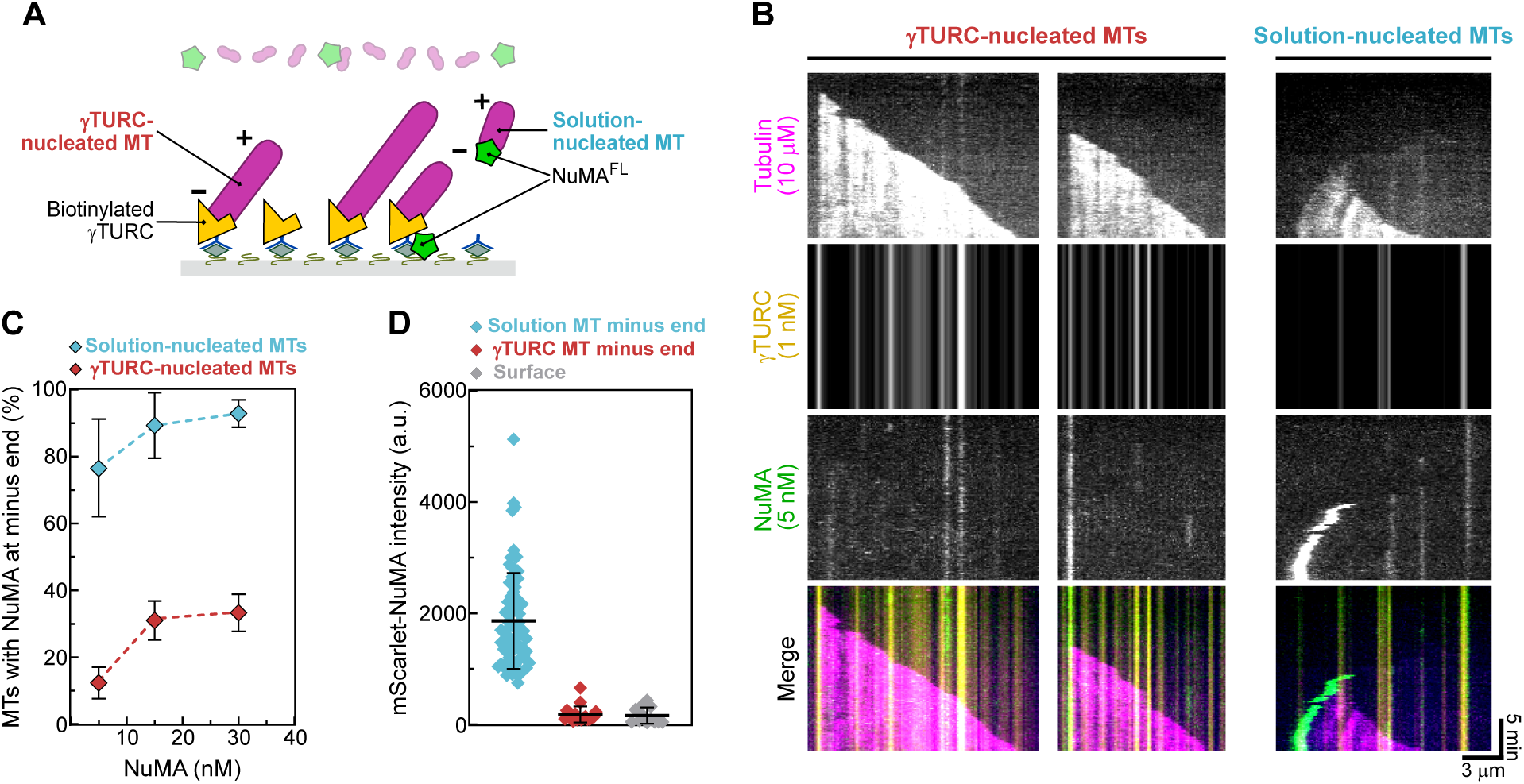
NuMA does not bind to γTuRC-capped microtubules. **(A)** Schematic of a γTURC nucleation assay performed in the presence of mScarlet-NuMA^FL^. **(B)** Representative TIRF microscopy kymographs showing microtubules nucleated by surface-immobilized, biotinylated mBFP-γTURC and spontaneously nucleated microtubules in solution in the presence of 10 µM Atto647-tubulin and 5 nM mScarlet-NuMA^FL^. γTURC and solution-nucleated microtubules can be distinguished because the latter diffuse on the surface. γTURC-nucleated microtubules occasionally display weak NuMA fluorescence at their minus ends (center), whereas the minus ends of solution-nucleated microtubules frequently show intense NuMA signal (right). **(C)** Frequency of apparent microtubule minus end localization of NuMA on γTURC and spontaneously nucleated microtubules in the presence of 10 µM Atto647-tubulin and 5 nM, 15 nM, 30 nM mScarlet-NuMA^FL^ (mean ± SEM, three independent experiments); bottom *n* = 80, 101, 104 microtubules; top *n* = 21, 54, 52 microtubules; apparent µTuRC-minus end co-localization frequency is not much higher than NuMA’s random surface adsorption (Methods). **(D)** Background-corrected maximum intensity of mScarlet-NuMA^FL^ at 30 nM localizing to either γTURC-nucleated microtubule minus ends, spontaneously nucleated microtubule minus ends, or at randomly chosen microtubule-free surface areas (mean ± SEM, three independent experiments), in the presence of 10 µM Atto647-tubulin. Each diamond represents the intensity at one microtubule minus end or surface area; *n* = 68, 18, 12.

γTuRC-nucleated microtubules grew only with their plus end, while their minus end was anchored to surface-immobilized γTuRC. In contrast, spontaneously nucleated microtubules in solution suddenly "landed" on the surface and moved diffusively. At the highest NuMA concentration tested, NuMA could be detected at the minus end of ≈93% of these spontaneously nucleated microtubules (Fig. 5 B, right, Fig. 5 C), whereas only ≈33% of the γTuRC-nucleated microtubules showed apparent localization of NuMA at the minus end, in most cases co-localizing with γTURC on the surface before microtubule nucleation (Fig. 5 B, middle, Fig. 5 C). The frequency of this apparent co-localization was not much higher than the estimated frequency of NuMA’s random surface adsorption (see Methods). In line with this, the fluorescence intensity of NuMA was much lower at γTURC-nucleated compared to solution-nucleated microtubule minus ends, and rather similar to the intensity of NuMA adsorbed to random microtubule-free surface areas (Fig. 5 D). Moreover, we observed that NuMA stimulated spontaneous nucleation of microtubules in solution in a dose-dependent manner, probably by stabilizing minus ends (Fig. S6 A). In contrast, no clear effect of NuMA on the efficiency of nucleation of microtubules by γTuRC could be observed (Fig. S6 B), in agreement with NuMA not interacting specifically with γTuRC-nucleated minus ends (Fig. 5 B and C).

In summary, the presence of γTuRC at microtubule minus ends hinders minus end binding of NuMA.

### NuMA binds to the minus ends of freshly severed microtubules

In cells, microtubules can be severed by severing enzymes, producing new minus ends (McNally and Roll-Mecak, 2018). Therefore, we tested whether NuMA could bind to new microtubule minus ends generated by microtubule severing using laser ablation (Fig. 6 A). After laser cutting of dynamic microtubules growing from surface-immobilized GMPCPP-seeds, NuMA indeed bound to and accumulated at the newly generated minus ends (Fig. 6 B and C). When the cut was executed close to the microtubule minus end, the severed short microtubule minus segment detached from the surface, diffusing away, and binding to the new minus end could be observed (Fig. 6 B). Cuts performed farther from the microtubule end produced longer microtubule segments that remained close to the surface, allowing to observe both freshly generated plus and minus ends (Fig. 6 C, Video 2). NuMA was never observed to bind to new plus ends, whereas ≈83% of the ablation-generated minus ends had NuMA bound (Fig. 6 D).

**Fig. 6.**
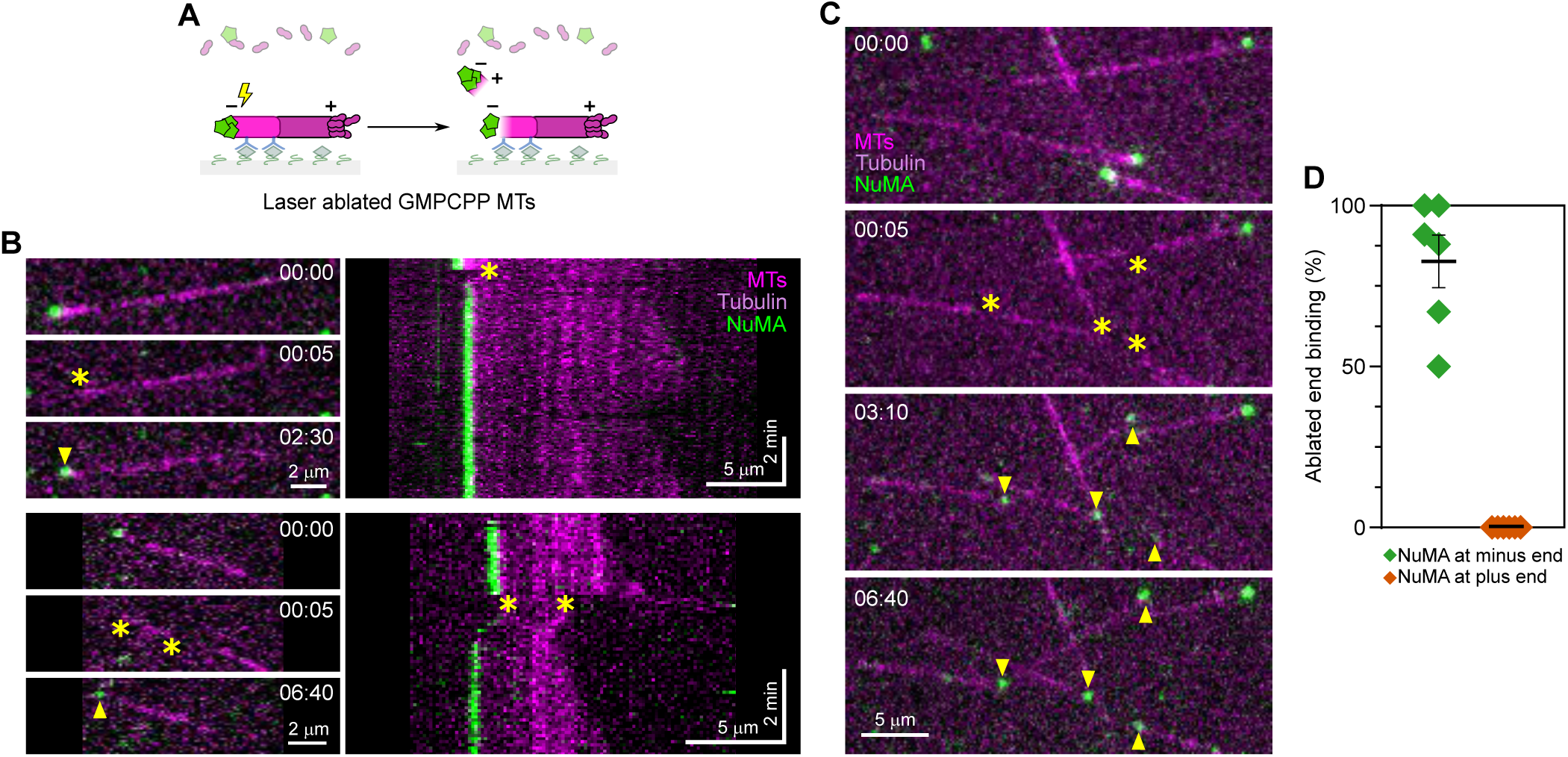
NuMA binds to the minus ends of freshly severed microtubules. **(A)** Schematic of a TIRF microscopy assay with microtubules elongating from surface-immobilized GMPCPP-seeds being severed by laser ablation in the presence of mScarlet-NuMA^FL^. **(B, C)** Representative TIRF microscopy time course images of a microtubule elongating from a biotinylated Atto647N-labelled GMPCPP-seed (magenta) in the presence of 40 nM mScarlet-NuMA^FL^ (green) and 10 µM Atto647N-tubulin (magenta). Upon severing of the microtubule by laser ablation (asterisks), NuMA binds selectively to the newly generated minus end (arrowheads). B: Ablation performed at one (top panel) or two (bottom panel) sites close to the microtubule minus end generates short segments that diffuse away in solution, thus only the minus end of the longest fragment which stays anchored to the surface can be observed. C: ablation performed farther from the microtubule minus end produces longer fragments that remain attached to the surface, allowing visualization of both new plus and minus ends. Related to Video 2. **(D)** Frequency of NuMA localization to freshly generated plus and minus ends after laser ablation in the presence of 40 nM mScarlet-NuMA^FL^ and 10 µM Atto647N-tubulin (mean ± SEM, three independent experiments). Each diamond represents an individual cycle of ablations followed by flowing of mScarlet-NuMA^FL^; multiple cycles were performed in each experiment); *n* = 59 ablations.

These data suggest that microtubule minus ends either released from γTuRC or produced by severing enzymes may be a suitable NuMA substrate in cells.

### Full-length NuMA can mediate dynein/dynactin-driven microtubule transport

We have demonstrated that full-length NuMA activates dynein motility via its N-terminal part and binds to and stabilizes free microtubule minus ends via its C-terminal part. This raises the question whether dynein together with full-length NuMA and dynactin can transport a microtubule minus end towards the minus end of another microtubule.

To test this, we allowed microtubules to nucleate and grow in suspension in the presence of NuMA^FL^ and GMPCPP, generating short stabilized microtubules with a strong accumulation of NuMA^FL^ at their minus ends (NuMA "lollipops") (Fig. 7 A). These NuMA lollipops were then added to immobilized long GMPCPP-microtubules that had been pre-incubated with dynein, dynactin and Lis1 (Fig. 7 B). Lollipops landed sometimes with their NuMA-decorated minus ends on the immobilized microtubules, showing then different types of behavior: ≈37% of lollipops could be observed to be transported processively towards the minus end of the immobilized microtubule, with NuMA and dynein bound to the lollipop minus end (Fig. 7 C, D and E). About half of these transport events showed a "dangling" lollipop with its minus end bound via dynein and NuMA to the immobilized microtubule (Fig. 7 C, D and E, Video 3). In a minority of cases, the lollipop aligned in a parallel fashion to the immobilized microtubule, apparently forming additional crosslinks, in addition to the dynein/NuMA link at the minus end of the lollipop microtubule (Fig. 7 E, Fig. S7 A and B). A larger fraction of transported lollipops was aligned to the immobilized microtubule in an antiparallel fashion, with their minus ends facing the plus end of the immobilized microtubule (Fig. 7 E, Fig. S7 C and D).

**Fig. 7.**
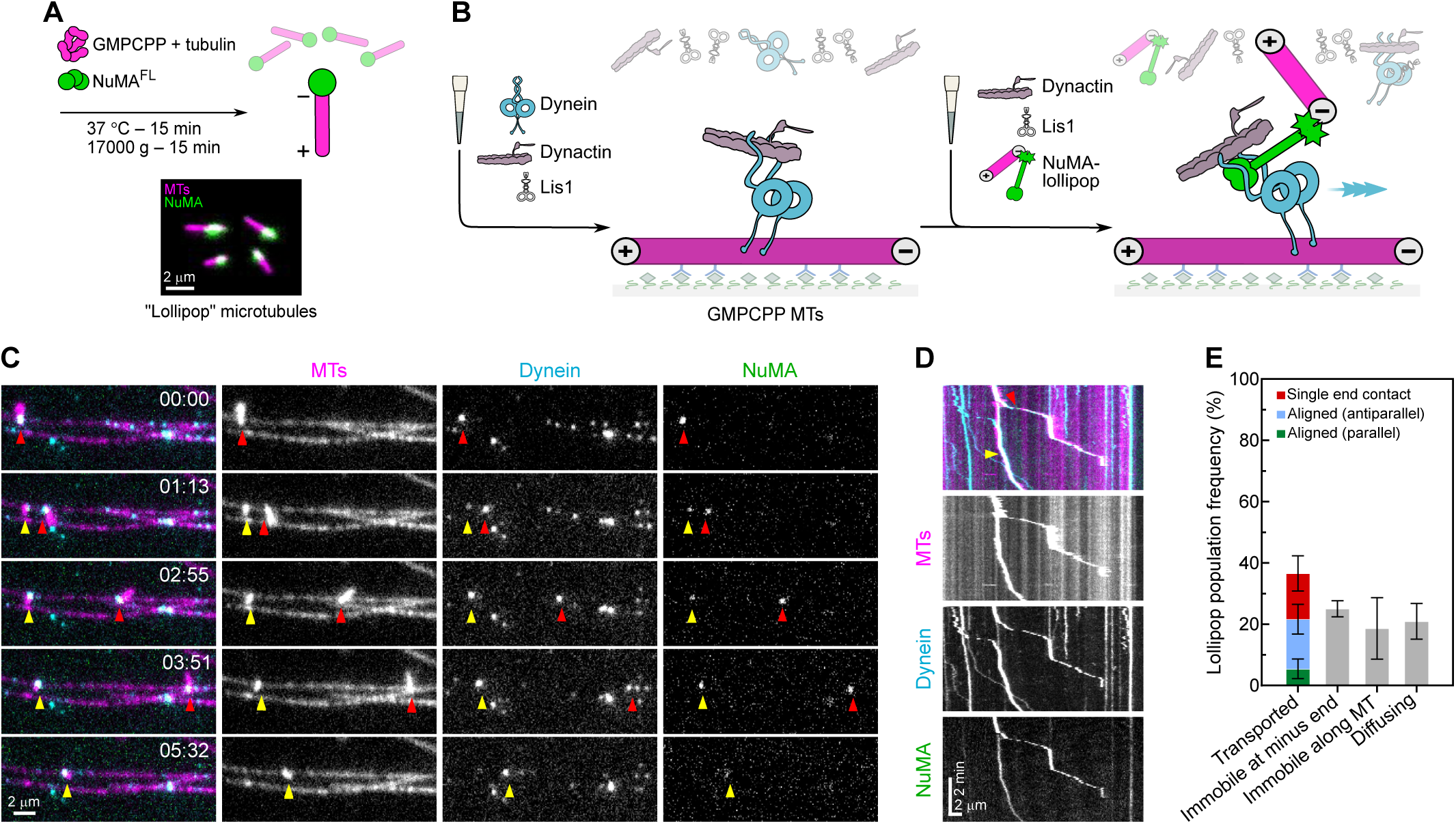
Full-length NuMA can mediate dynein/dynactin-driven microtubule transport. **(A)** Schematic (top) of the polymerization protocol to obtain microtubule “lollipops”, with representative TIRF microscopy image (bottom) of lollipops obtained from a mixture of 0.8 µM Atto647N-tubulin (magenta), 40 nM mScarlet-NuMA^FL^ (green) and 1 mM GMPCPP. Minus ends can be identified by the selective presence of mScarlet-NuMA^FL^. **(B)** Schematic of a two-flush TIRF microscopy dynein-driven microtubule transport assay: first, dynein, dynactin and Lis1 are flowed into a channel containing long immobilized GMPCPP-microtubules; dynein is allowed to accumulate on microtubules for ≈3 min; in a second step, lollipops are introduced into the channel, with dynactin and Lis1, leading to dynein/dynactin/lollipop-bound NuMA transporting lollipop microtubules. **(C, D)** Representative time course TIRF microscopy images (C) and related kymograph (D) of transport events performed in the presence of 14 nM mEGFP-dynein (pre-bound to immobilized GMPCPP-microtubules), 28 nM dynactin and 1000 nM Lis1. For both the faster (red arrowheads) and slower (yellow arrowheads) transport events, the lollipop is bound to the immobilized microtubule through a single anchoring point, corresponding to co-localizing mEGFP-dynein and mScarlet-NuMA^FL^ (arrowheads), resulting in it to dangle while being transported. Related to Video 3. **(E)** Frequency of lollipop microtubule behaviors on immobilized microtubules (mean ± SEM, three independent experiments); protein concentrations as in C‒D; *n* = 82 landed lollipops.

≈25% of the lollipops were bound in a static manner with their minus ends at the minus end of the immobilized microtubules (Fig. 7 E), a configuration that might have resulted from transport before imaging started. Alternatively, lollipops might have landed directly on the dynein-decorated minus ends of the immobilized microtubules (Fig. S7 E).

Lollipop microtubule transport velocities were varied, ranging from 3 nm s^-1^ to 400 nm s^-1^, considerably slower than the velocities of the dynein/dynactin/NuMA^FL^ complex alone (Fig. S2 B). This may be due to friction generated by some immobile dynein molecules that are typically observed under such *in vitro* conditions, or by NuMA clusters that may generate immobile crosslinks between microtubules.

Taken together, these observations demonstrate that NuMA is a dynein adaptor that can bind microtubule minus ends as a cargo, thus enabling the dynein-driven gathering of minus ends.

## DISCUSSION

Using *in vitro* reconstitution experiments, we have established NuMA as the first mitotic activating dynein adaptor, and extended the list of categorized dynein cargoes, now including also microtubule minus ends. Our results support a model for spindle pole focusing where NuMA activates dynein/dynactin via its N-terminus and at the same time connects it to a microtubule minus ends via its C-terminus, allowing minus ends to be transported to other microtubule minus ends.

We found that NuMA’s N-terminal part is sufficient to act as activating dynein adaptor, but compared to other previously studied adaptors (McKenney et al., 2014; Schlager et al., 2014; Schroeder and Vale, 2016; Redwine et al., 2017; Urnavicius et al., 2018; Canty et al., 2023), efficient initiation of dynein’s processive motility by NuMA in the presence of dynactin is strictly dependent on the additional presence of Lis1, which is known to relieve dynein’s auto-inhibition and thereby promote dynein/dynactin/adaptor formation (Qiu, Zhang and Xiang, 2019; Elshenawy et al., 2020; Htet et al., 2020; Marzo, Griswold and Markus, 2020; Karasmanis et al., 2023). This additional dependence may suggest a different binding mode of NuMA to dynein/dynactin, in agreement with differences in the arrangement of known adaptor motifs (Olenick and Holzbaur, 2019). NuMA contains both a Hook domain and a CC1-box-like motif (Renna et al., 2020), a unique combination among established dynein adaptors. Our results with purified proteins agree well with studies in cells and *Xenopus* egg extract showing that dynein, dynactin, NuMA and Lis1 are all needed for correct spindle pole focusing (Wang et al., 2013; Monda and Cheeseman, 2018; So et al., 2022).

Going beyond the previously known microtubule-binding capacity of NuMA’s C-terminal part, we discovered here that NuMA binds directly and preferentially to microtubule minus ends, stopping their growth and protecting them against shrinkage, effectively capping them. The observation that γTuRC hinders NuMA’s minus-end binding may suggests that NuMA recognizes either the exposed α-tubulins at minus ends or even the interior of microtubules at their minus ends, both being made inaccessible by γTuRC (Aher et al., 2024; Brito et al., 2024; Dendooven et al., 2024). Structural studies will be needed to provide more insight into the mechanism of microtubule minus end capping by NuMA.

The importance of both the N-terminal and C-terminal part being required for NuMA’s function to contribute to pole focusing is in agreement with experiments in cells or *Xenopus* egg extract showing that depleting NuMA (Merdes et al., 1996, 2000; Hueschen et al., 2017; Sun et al., 2021; Toorn et al., 2022), removing, over-expressing or mutating its N-terminal part (Kotak, Busso and Gönczy, 2012; Okumura et al., 2018; Renna et al., 2020), or removing its C-terminal part (Silk, Holland and Cleveland, 2009; Hueschen et al., 2017; Okumura et al., 2018) leads to strong spindle pole focusing defects. However, some reported results appear to contradict our simple model, especially when only parts of NuMA’s tail were removed, potentially suggesting that other NuMA interactors or post-translational modifications not present in our reconstitutions may contribute to NuMA’s pole focusing function (Hueschen et al., 2017; Okumura et al., 2018). NuMA is the only activating dynein adaptor known to date that connects dynein to a cytoskeletal filament. Linking a microtubule-dependent motor to a microtubule minus end generates a new type of active microtubule crosslinker. Dynein/dynactin/NuMA is different from the plus-end directed motor kinesin-5 (KIF11 in human) or kinesin-12 (KIF15 in human) that are symmetric crosslinkers with dimeric motors at both ends of the molecule, allowing them to move along two microtubule simultaneously (Kashina et al., 1996; Kapitein et al., 2005; Drechsler and McAinsh, 2016). The minus-end directed kinesin-14 (HSET or KIFC1 in human) is an asymmetric crosslinker that links two microtubules using a dimeric motor connected to a diffusive microtubule binding domain without any microtubule end specificity(Fink et al., 2009; Hentrich and Surrey, 2010). This design, however, causes kinesin-14 to become a poor pole focusing motor in the presence of kinesin-5 (Henkin et al., 2022). Similar to kinesin-14, dynein/dynactin/NuMA is also an asymmetric microtubule crosslinker with one motor end and a non-motor microtubule-binding end, however the latter binds microtubules with a strong preference for minus ends. This design appears to be specialized for efficient minus-end focusing, suggesting that different motor designs have evolved for distinct functions. In the future, it will be interesting to understand how this new type of crosslinker cooperates with other microtubule crosslinking motors of different designs to ensure correct bipolar spindle organization.

Our observation that γTuRC at microtubule minus ends prevents NuMA minus-end binding raises an important question: how can minus ends of freshly γTuRC-nucleated microtubules in the spindle be brought together by dynein/dynactin/NuMA? Given that we could show that NuMA can bind to freshly severed minus ends, it appears that minus ends need to be uncapped from γTuRC or microtubules need to be severed before free minus ends can then be bound by NuMA and transported to the spindle pole by dynein. This model agrees with the observed enrichment of microtubule severing enzymes on spindles, especially near the poles (Jiang et al., 2017; Jin et al., 2017), and with the observed NuMA accumulation at the minus ends of kinetochore fibers after laser ablation in cells (Hueschen et al., 2017). The requirement for γTuRC uncapping may prevent excessive concentration of γTuRC at spindle poles, given that it also needs to be available throughout the spindle for augmin/RanGTP-dependent branched microtubule nucleation(Uehara et al., 2009; Petry et al., 2013). On the other hand, when being concentrated at poles by dynein transport, NuMA’s reported clustering and condensation activity may then further stabilize spindle poles (Okumura et al., 2018; Sun et al., 2021).

NuMA is also known to bind the membrane-anchored protein LGN with its C-terminal tail, thereby tethering dynein to the cortex and allowing dynein to position the spindle correctly in the cell (Okumura et al., 2018; Pirovano et al., 2019). Phosphorylation by the mitotic kinases CDK1 (Compton and Luo, 1995; Kotak, Busso and Gönczy, 2013; Seldin et al., 2013), Aurora-A kinase (Gallini et al., 2016; Kotak et al., 2016) and Polo-like kinase 1 (Plk1) (Kettenbach et al., 2011; Sana et al., 2018) have been shown to control the relative distribution and turnover of NuMA at the spindle poles and the cell cortex. In the future, it will be important to understand how these phosphorylations may selectively affect NuMA’s interaction with either cortical binding sites versus microtubule minus ends at the spindle poles, or with dynein/dynactin or other potential binding partners to achieve the right balance of dynein/NuMA activity at different cellular locations and times during mitosis.

## METHODS

All reagents were purchased from Sigma-Aldrich, unless otherwise stated. Chromatography columns were purchased from Cytiva.

### DNA constructs

All NuMA constructs were generated from a plasmid, containing the coding sequence for full-length human NuMA isoform 1 (NM_006185), codon-optimized for *Spodoptera frugiperda* expression (generous gift of Marina Mapelli). All vectors shared the same pFastBac1 (Invitrogen) backbone. NuMA^N-term^ was N-terminally tagged with sequences encoding for a Strep-tag II, a tobacco etch virus (TEV) protease cleavage site, a SNAP-tag, and PreScission protease cleavage site. All other NuMA constructs encoded for an mScarlet instead of a SNAP-tagand lacked the PreScission site. To generate NuMA^N-term^, two consecutive stop codons were inserted after the coding sequence for the first 705 aa of full-length NuMA. To create NuMA^C-term^ ^L^, NuMA^C-term^ ^S1^ and NuMA^C-term^ ^S2^, the coding sequences for aa 1560‒2115, 1701‒1981 and 2002‒2105, respectively, were amplified from the full-length sequence. Constructs containing the C-terminal part of NuMA have D1560T, T1820A, H2115InsLE changes compared to the sequence in the database.

To produce a construct for the simultaneous expression in insect cells of all six subunits of the human cytoplasmic dynein 1 complex, the biGBac system was utilized (Weissmann et al., 2016). First, we amplified the sequence encoding for an N-terminal His_8_-tag followed by a ZZ-tag, two TEV protease cleavage sites, an mEGFP, and the dynein heavy chain (DHC) (*DYNC1H1*, NM_001376.4), from the vector pGFPdyn1 (Jha et al., 2017), and cloned it into the pBIG1a vector. Compared to the canonical EGFP sequence, our gene contains two changes: i) M1insS, resulting from the cloning process; ii) L221K, previously shown to promote a monomeric state (Zacharias et al., 2002; Snapp et al., 2003). Secondly, each of the sequences encoding for the smaller subunits intermediate chain 2C (IC2C) (*DYNC1I2*, AF134477), light intermediate chain 2 (LIC2) (*DYNC1LI2*, NM_006141.2), Tctex1 (*DYNLT1*, NM_006519.2), roadblock 1 (Robl1) (*DYNLRB1*, NM_014183.3) and light chain 8-1 (LC8-1) (*DYNLL1*, NM_003746.2) was amplified from a pDyn2 vector (Schlager et al., 2014) (generous gift of Andrew Carter) and separately cloned into the pLIB vectors. The individual sequences were then amplified from their respective pLIB and cloned together into the pBIG1b vector. Eventually, all dynein subunits were joined into the pBIG2ab vector.

The coding sequence for full-length human Lis1 was amplified from the plasmid His_6_-TEV-mCherry-Lis1-pFastBac1 (Jha et al., 2017) and cloned into the vector pFastBacHT A (Invitrogen) to obtain a construct for insect cell expression of unlabeled Lis1 N-terminally tagged with an His_6_-tag and TEV protease cleavage site.

To generate a construct for bacterial expression of BicD2N^1‒400^, the coding sequence for the first 400 aa of human BicD2 was amplified from a cDNA (Origene, SC300552) by PCR and cloned into a pETZT2 plasmid, inserting N-terminally a His_6_-tag, a Z-tag, and TEV cleavage site.

Lentiviral vectors for the expression of a fluorescently-tagged and biotinylatable human γTuRC were generated as detailed previously (Consolati et al., 2020).

### Cell lines

*Escherichia coli* strains DH5α, BL21-CodonPlus (DE3)-RIPL, and MAX Efficiency DH10Bac (Gibco) were grown in Luria Bertani (LB) medium (CRG Protein Technologies Unit) in the presence of appropriate antibiotics. For expression of recombinant proteins in insect cells, the *Spodoptera frugiperda* strain Sf21 (source: EMBL) was grown in suspension at 27 °C in Sf-900TM III SFM Serum Free Medium (Gibco). The baculovirus preparation was carried out according to the manufacturer’s protocol (Bac-to-Bac system, Life Technologies), and baculovirus-infected insect cells were frozen to generate stable viral stocks similarly to what described previously (Wasilko et al., 2009). For recombinant biotinylated mBFP-γTURC expression, HeLa-Kyoto cells (RRID:CVCL_1922) were cultured, infected and harvested as detailed previously (Consolati et al., 2020).

### Purification and labeling of recombinant human mScarlet-NuMA^N-term^

Recombinant human NuMA^N-term^ was purified from a pellet of 1.6 L of Sf21 cell culture (≈16 g) resuspended in lysis buffer (50 mM sodium phosphate buffer, 300 mM KCl, 0.5 mM adenosine-5’-triphosphate [ATP], 5 mM 2-mercaptoethanl [2-ME], pH 7.4) supplemented with 50 U ml^-1^ Benzonase and protease inhibitors. The lysis was carried out using an Avestin EmulsiFlex-C5 homogenizer (two rounds). After clarification of the lysate by centrifugation (15,800 × *g*, 30 min, 4 °C), the supernatant was loaded onto a 5 ml StrepTrap HP column. The column was washed with lysis buffer, and the elution was carried out using lysis buffer supplemented with 2.5 mM D-desthiobiotin. The NuMA-containing fractions were pooled, concentrated (Amicon Ultra 4, 30 kDa MWCO), and the Strep-tag II was cleaved off by incubating with TEV protease (CRG Protein Technology Unit) for 2 h at 4 °C, using a TEV-to-protein ratio of 1:15 (wt/wt). During or after cleavage, 1.5 ml fractions of the protein pool were incubated with 3-fold molar excess of SNAP-Surface Alexa Fluor 546 or SNAP-Surface Alexa Fluor 647 (New England Biolabs) overnight at 4 °C. The labelled protein was concentrated (Amicon Ultra 15, 50 kDa MWCO) and ultra-centrifuged (278,088 × *g*, 10 min, 4 °C). The unreacted dye was removed by size exclusion chromatography using a Superose 6 Increase 10/300 GL column and lysis buffer. The NuMA-containing peak fractions were pooled and concentrated to ≈0.4 mg ml^-1^. The yield was ≈0.2 mg protein/g pellet and the labelling efficiency 95‒100%.

### Purification of recombinant human mScarlet-NuMA^FL^

Recombinant human mScarlet-NuMA^FL^ was purified from a pellet of 2.1 L of Sf21 cell culture (≈15 g) resuspended in lysis buffer (50 mM sodium phosphate buffer, 150 mM KCl, 1 mM EDTA ethylenediaminetetraacetic acid [EDTA], 2 mM MgCl_2_, 0.5 mM ATP, 0.02% Brij-35 [ThermoFisher Scientific], 5 mM 2-ME, pH 8.0) supplemented with 50 U ml^-1^ Benzonase (Merck-Millipore), 50 µg ml^-1^ DNAse I (Roche Applied Sciences) and protease inhibitors (cOmplete EDTA-free Protease Inhibitor Cocktail [Roche Applied Science]). Lysis was carried out using a douncer homogenizer (Wheaton, 50 tight strokes). The lysate was supplemented with 350 mM KCl and incubated 10 min on ice. After clarification by centrifugation (256,631 × *g*, 45 min, 4 °C), the supernatant was loaded onto a 5 ml StrepTrap HP column, equilibrated with binding buffer (lysis buffer supplemented with 350 mM KCl and protease inhibitors). The column was washed with wash buffer (lysis buffer supplemented with 350 mM KCl and 4.5 mM ATP), and the protein was eluted using binding buffer supplemented with 10 mM D-desthiobiotin. The NuMA-containing fractions were pooled and concentrated (Amicon Ultra 4, 100 kDa MWCO [Merck Millipore]). The Strep-tag II was cleaved off by incubating with TEV protease for 2 h at 4 °C, using a TEV-to-protein ratio of 1:10 (wt/wt), and the digested protein was ultra-centrifuged (278,088 × *g*, 10 min, 4 °C). The protein was further purified by size exclusion chromatography using a Superose 6 Increase 10/300 GL column with size exclusion buffer (50 mM sodium phosphate buffer, 300 mM KCl, 0.2% Brij-35, 2 mM DTT [NZYtech], pH 8.0) supplemented by protease inhibitors. The NuMA-containing peak fractions were pooled and concentrated to ≈1.5 mg ml^-1^. The yield was ≈0.3 mg protein/g pellet.

### Purification of recombinant human mScarlet-NuMA^C-term L^

Recombinant human NuMA^C-term^ ^L^ was purified from a pellet of 1.4 L of Sf21 cell culture (≈10 g), resuspended in lysis buffer (50 mM sodium phosphate buffer, 500 mM KCl, 0.5 mM ATP, 5 mM 2-ME, pH 7.8) supplemented with 120 U ml^-1^ Benzonase, 2 mM phenylmethylsulfonyl fluoride (PMSF), and protease inhibitors. The purification protocol resembled that of NuMA^N-term^, with the following differences: i) the speed of the first centrifugation was 20,400 × *g*; ii) after a first wash with lysis buffer supplemented with protease inhibitors, the StrepTrap HP column was washed with lysis buffer supplemented by 500 mM KCl; iii) the NuMA-containing StrepTrap HP eluate fractions were not concentrated prior to cleavage by TEV. The final pool of NuMA-containing peak fractions were concentrated to ≈0.6 mg ml^-1^. The yield was ≈0.5 mg protein/g pellet.

### Purification of recombinant human mScarlet-NuMA^C-term S1^

Recombinant human NuMA^C-term^ ^S1^ was purified from a pellet of 1 L of Sf21 cell culture (≈11 g) resuspended in lysis buffer (50 mM KH_2_PO_4_, 50 mM Na_2_HPO_4_ [ThermoFisher Scientific], 800 mM KCl, 2 mM MgCl_2,_ 1 mM EDTA, 2 mM 2-ME, pH 8.0) supplemented with 50 U ml^-1^ Benzonase and protease inhibitors. Lysis was carried out using an Avestin EmulsiFlex-C5 homogenizer (two rounds). After clarification of the lysate by centrifugation (30,000 × *g*, 30 min, 4 °C), the supernatant was filtered through a Millex-HV sterile syringe filter unit PVDF (0.45 μm, Merck), and loaded onto a 5 ml StrepTrap HP column. The column was washed with lysis buffer and eluted with Strep elution buffer (50 mM KH_2_PO_4_, 50 mM Na_2_HPO_4_, 500 mM KCl, 2 mM MgCl_2,_ 1 mM EDTA, 2 mM 2-ME, 2.5 mM D-desthiobiotin, pH 8.0). The NuMA-containing fractions were pooled and concentrated (Amicon Ultra 4, 30 kDa MWCO). The Strep-tag II was cleaved off by incubating with TEV protease overnight at 4 °C, using a TEV-to-protein ratio of 1:30 (wt/wt). The cleaved protein was concentrated (Amicon Ultra 4, 30 kDa MWCO), ultra-centrifuged (529,484 × *g*, 10 min, 4 °C), and loaded on a HiLoad 16/600 Superdex 200 prep grade column. The elution was performed using size exclusion buffer (BRB80 (80 mM K-PIPES, 1 mM MgCl_2,_ 1 mM ethylene glycol bis(β aminoethyl ether) N,N,N’,N’ tetraacetic acid [EGTA]), 500 mM KCl, 5 mM 2-ME, pH 6.8). The NuMA-containing peak fractions were pooled, exchanged into storage buffer (BRB80, 50 mM KCl, 5 mM 2-ME, pH 6.8), and concentrated (Amicon Ultra 4, 30 kDa MWCO) to ≈15 mg ml^-1^. The yield was ≈1 mg protein/g pellet.

### Purification of recombinant human mScarlet-NuMA^C-term S2^

Recombinant human NuMA^C-term^ ^S2^ was purified from a pellet of 0.9 L of Sf21 cell culture (≈4 g) using the same protocol described for NuMA^C-term^ ^S1^ with two modifications: i) prior to TEV cleavage, the protein was exchanged into cleavage buffer (50 mM KH_2_PO_4_, 50 mM Na_2_HPO_4_, 200 mM KCl, 2 mM MgCl_2,_ 1 mM EDTA, 2 mM 2-ME, pH 7.0) using a PD-10 desalting column; ii) the size exclusion elution was carried out in storage buffer, which avoided the final buffer exchange step. The protein was concentrated to ≈24 mg ml^-1^ and the yield was ≈4 mg protein/g pellet.

### Purification of recombinant human mEGFP-dynein complex

Recombinant human mEGFP-dynein was purified from a pellet of 1.4 L of Sf21 cell culture (≈16 g) as described (Jha et al., 2017) with the following modifications: i) 250 mM K-acetate in the lysis buffer was substituted by 150 mM KCl; ii) the lysate was clarified twice by ultra-centrifugation (125,749 g, 20 min, 4 °C; 225,634 × *g*, 20 min, 4 °C); iii) size exclusion chromatography was performed using a HiLoad 16/600 Superose 6 prep grade column with size exclusion buffer (50 mM HEPES, 400 mM K-acetate, 2 mM MgSO_4_, 0.1 mM ATP, 5 mM DTT, pH 7.4). The dynein-containing peak fractions were concentrated (Amicon Ultra 4, 50 kDa MWCO) to ≈1.2 mg ml^-1^. The yield was ≈0.3 mg protein/g pellet.

### Purification of porcine brain dynactin

Endogenous brain dynactin was purified from two pig brains by BicD2N^1‒400^ affinity chromatography followed by ion exchange chromatography, as described (Jha et al., 2017), with the following modifications: i) to generate the BicD2N^1‒400^ column, 60 mg of purified His_6_-tag-Z-tag-TEV-BicD2N^1‒400^ (see below) were conjugated to a 5 ml HiTrap NHS-Activated HP column following the manufacturer’s recommended method, with a ≈90% yield; ii) two intermediate steps of washing with lysis buffer were carried out during the application of the lysate to the BicD2 column to avoid overloading and clogging; iii) the dynein-dynactin complex was eluted at 3 ml min-1 using lysis buffer supplemented with 1 M KCl through a 25 ml step gradient from 0 to 500 mM KCl; iv) the flow-through was collected and re-loaded onto the BicD2N^1‒400^ column to recover leftover unbound dynein-dynactin complex of the first loading, for a maximum of two times; v) for the elution of the MonoQ 5/50 GL column, the following gradient was applied, using MonoQ binding buffer supplemented with 800 mM NaCl: linear 0‒224 mM in 10 ml; step 224‒256 mM in 1 ml; linear 256‒272 mM in 20 ml; step 272‒312 mM in 1 ml; linear 312-400 mM in 10 ml. The dynactin complex eluted at 330 mM NaCl. The dynactin-containing peak fractions were concentrated (Amicon Ultra 4, 50 kDa MWCO) to ≈0.6 mg ml^-1^. The yield was ≈75 µg protein/g pellet.

### Purification of recombinant human Lis1 constructs

Recombinant human mCherry-Lis1 was purified according to a published protocol (Jha et al., 2017). Unlabeled Lis1 was purified from a pellet of 0.8 L of Sf21 cell culture (≈ 11 g) resuspended in lysis buffer (50 mM sodium phosphate, 500 mM NaCl, 20 mM imidazole, 2 mM MgCl_2_, 10% glycerol [vol/vol, Fisher Scientific], 0.5 mM ATP, 5 mM 2-ME, pH 7.4) supplemented with 20 µg ml^-1^ DNAse I, protease inhibitors, and 0.5% Triton X-100 (vol/vol). Cells were lysed using a douncer homogenizer (50 tight strokes). The lysate was clarified by centrifugation (104,350 × *g*, 15 min, 4 °C) and run on a 5 ml HisTrap FF column. The column was washed first with lysis buffer supplemented with 500 mM NaCl, 4.5 mM ATP, and 8 mM MgCl_2_; secondly with two step gradients using lysis buffer supplemented with 330 mM imidazole: i) 20‒27.5 mM imidazole; ii) 27.5‒55 mM imidazole. The protein was eluted using lysis buffer supplemented with 330 mM imidazole. The protein-containing fractions were pooled and exchanged into size-exclusion buffer (50 mM HEPES, 300 mM KCl, 0.05 mM ATP, 10% glycerol [vol/vol], 2 mM DTT, pH 7.4) using a HiTrap desalting column. The His_6_-tag was cleaved off by overnight incubation with TEV protease at 4 °C, using a TEV-to-protein ratio of 1:50 (wt/wt). The cleaved protein was concentrated (Amicon Ultra 15, 30 kDa MWCO), ultra-centrifuged (278,088 × *g*, 10 min, 4 °C), and loaded onto a Superdex 200 10/300 GL column. The elution was performed with size-exclusion buffer; the Lis1-containing peak fractions were pooled and concentrated (Amicon Ultra 4, 30 kDa MWCO) to ≈9 mg ml^-1^. The yield was ≈0.3 mg protein/g pellet.

### Purification of recombinant human BicD2N^1‒400^

Recombinant human BicD2N^1‒400^ was expressed in *E. coli* BL21-CodonPlus (DE3)-RIPL cells by induction with 0.5 mM IPTG for 16 h at 16 °C. A pellet of 1 L of culture (≈4 g) was resuspended in lysis buffer (50 mM HEPES, 400 mM KCl, 2 mM MgCl_2_, 0.1 mM ATP, 10 mM imidazole, 2 mM 2-ME, pH 7.4) supplemented with 2 mM PMSF and protease inhibitors. Lysis was carried out using an Avestin EmulsiFlex-C5 homogenizer (two rounds). The lysate was clarified by centrifugation (256,631 × *g*, 20 min, 4 °C) and run over two 5 ml HisTrap HP columns. The columns were washed with wash buffer (50 mM HEPES, 1 M KCl, 2 mM MgCl_2_, 2 mM ATP, 2 mM 2-ME, pH 7.4), and the elution was performed with lysis buffer supplemented with 490 mM imidazole. For each column, the flow-through was re-loaded to recover the leftover unbound BicD2N^1‒400^. The protein-containing fractions were pooled and the His_6_-tag was cleaved off by overnight incubation with TEV protease at 4 °C, using a TEV-to-protein ratio of 1:30 (wt/wt). The cleaved protein was concentrated (Amicon Ultra 15, 30 kDa MWCO), ultra-centrifuged (529,484 × *g*, 10 min, 4 °C, 15 min, 4 °C), and loaded onto four HiLoad 16/600 Superdex 200 prep grade columns. The elution was carried out using size exclusion buffer (50 mM HEPES, 200 mM KCl, 1 mM MgCl_2_, 10% glycerol [vol/vol], 1 mM 2-ME, pH 7.4). The pool of BicD2N^1‒400^-containing peak fractions was split in two parts: the one destined to NHS-coupling (see dynactin purification above) was kept at ≈3.4 mg ml^-1^; the one destined to microscopy assays was concentrated (Amicon Ultra 4, 30 kDa MWCO) to ≈12 mg ml^-1^. The yield was ≈5 mg protein/g pellet.

### Purification of recombinant human biotinylated mBFP-γTuRC

Recombinant human biotinylated mBFP-γTuRC was purified from HeLa-Kyoto cells as described previously (Brito et al., 2024).

### Purification and labeling of porcine brain tubulin

Endogenous tubulin was isolated from pig brain following sequential cycles of polymerization-depolymerization as previously described (Hyman, 1991; Castoldi and Popov, 2003). Tubulin was further purified by recycling and part of it was labeled with Atto647-NHS, Atto647N-NHS, and EZ-Link NHS-Biotin (ThermoFisher Scientific), according to published methods (Consolati et al., 2022).

### Chromatography and protein concentrations

All purification steps were carried out at 4 °C; all buffers were degassed and chilled to 4 °C prior to pH adjustment; all types of chromatography were performed using an ÄKTA Pure System. At the final stage of every purification, the protein-containing peak fractions of the size exclusion eluate were identified by sodium dodecyl sulfate polyacrylamide gel electrophoresis (SDS-PAGE) Coomassie Blue G-250 staining. The concentrated fraction pool was ultra-centrifuged (278,088 × *g*, 10 min, 4 °C), and flash-frozen prior to long-term storage in liquid nitrogen. The protein concentrations were calculated after freeze-thawing from the absorbance measured at 280 nm via Bradford assay (Protein Assay Dye Reagent Concentrate [Bio-Rad]). For NuMA^C-term^ ^L^, the concentration was derived by SDS-PAGE Coomassie Blue G-250 staining quantification. For Atto647N-tubulin, Atto647-tubulin, AF546-NuMA^N-term^, and AF647-NuMA^N-term^ the dye concentration was obtained from the absorbance measured at the dye-specific wavelength and the extinction coefficient of the dye. Protein concentrations refer to monomers, except for tubulin concentrations, which refer to heterodimers, and dynein and dynactin concentrations refer to one copy of the entire complex.

### SDS-PAGE and Western blotting

Protein samples were resolved by SDS-PAGE (Fig. S1 A, Fig. S3 A and B) using two electrophoresis systems: i) XCell SureLock Mini-Cell (Invitrogen), in combination with NuPAGE Bis-Tris Mini protein gels (Invitrogen), NuPAGE LDS Sample Buffer 4X (Invitrogen), NuPAGE MES and MOPS SDS running buffers 20X (Invitrogen); ii) Mini-PROTEAN Tetra Cell (Bio-Rad), in combination with Mini-PROTEAN TGX precast protein gels (Bio-Rad), Laemmli SDS sample buffer reducing 6X (Alfa Aesar), XT-Tricine running buffer (Bio-Rad). Precision Plus Protein Dual Xtra (Bio-Rad) and HiMark (Invitrogen) were used as pre-stained protein standard. Gels were run according to the manufacturers’ recommendations. Gel staining was performed using InstantBlue Coomassie protein stain (Abcam) or Coomassie Brilliant Blue R-250 dye (ThermoFisher Scientific).

For immunoblotting of dynein and dynactin subunits (Fig. S1 A), the gels were transferred to iBlot2 Transfer Stacks PVDF (0.2 µm, Invitrogen) using the iBlot 2 Gel Transfer Device (Invitrogen). Dynein heavy chain subunit was transferred at 25V for 12 min; dynein lighter chains using the default protocol “P3”; all dynactin subunits using the default protocol “P0”. Membrane blocking and antibody dilution buffer consisted of: tris-buffered Saline (TBS) or phosphate-buffered saline (PBS) (CRG Protein Technologies Unit), 5% skimmed milk powder (wt/vol, Millipore), 0.05% Tween20 (vol/vol). Anti-GFP, anti-dynein antibodies, anti-dynactin antibodies, and HRP-conjugated secondary antibodies were diluted according to the manufacturers’ recommendations. Stained gel imaging and blot chemiluminescent detection were carried out using iBright CL1500 Imaging System (Invitrogen). Fluorescence visualization of mScarlet-tagged NuMA bands (Fig. S3 B) was performed using Molecular Imager Gel Doc XR System (Bio-Rad).

### Mass photometry

The oligomerization state of all proteins introduced in this study, except for mScarlet-NuMA^FL^, was analyzed using a TwoMP mass photometer (Refeyn) (Fig. S1 B, Fig. S3 C). All measurements and dilutions were executed with the respective TIRF microscopy assay buffer; methylcellulose was however excluded from the buffer. For each sample, 18 µL of buffer was loaded onto a well (3 mm diameter, Refeyn) for the autofocus; 2 µl of protein dilution was then mixed into the buffer drop to reach a final concentration of 5‒20 nM. Protein contrast count was acquired with at least two technical replicates. Molecular weights were assigned by comparison with calibration probes of known mass (Native Mark unstained protein standard [Invitrogen], β-amylase). All data was processed using the DiscoverMP software (Refeyn).

### Stabilized microtubules

#### Stabilized microtubules for microtubule sedimentation assays

Microtubules for sedimentation assays were prepared from a mixture containing: 15 µM tubulin, including 16% (mol/mol) of Atto647N-tubulin (for a final fluorescent labeling ratio of 6.2%); 1 mM guanosine 5’ [(α,β) methyleno] triphosphate (GMPCPP) (Jena Bioscience); BRB80 to reach a final volume of 60 µl. The mixture was incubated for 5 min on ice and then 1 h at 37 °C. The microtubules were centrifuged in a tabletop centrifuge at 17,000 × *g* for 20 min; the pellet was resuspended in 100 µl BRB80T (BRB80 supplemented by 10 µM Paclitaxel), re-centrifuged at 17,000 × *g* for 10 min, finally resuspended in the same volume of BRB80T, and used within the same day.

#### Long stabilized microtubules for microscopy

To generate double stabilized long microtubules, we incubated 1.3 µM tubulin containing 43% (mol/mol) biotin-labeled tubulin and 10% (mol/mol) of Atto647N-tubulin (for a final fluorescent labeling ratio of 3.8%) in BRB80 with 1 mM GMPCPP, 1.25 mM MgCl_2_, and 1 mM tris(2 carboxyethyl)phosphine (TCEP) (90 µl final volume) for 5 min on ice, and then at 27°C overnight. Centrifugation was carried out as described above for *Stabilized microtubules for microtubule sedimentation assays,* and the pellet was resuspended in 90 µl BRB80T. These long, biotin and Atto647N-labelled microtubules were stored at room temperature and used for up to two weeks. Prior to every usage, they were centrifuged as described above. For microscopy assays, the microtubule preparation was diluted up to 5-fold in BRB80T.

#### Short stabilized microtubules (“seeds”) for microscopy

Stabilized short and bright GMPCPP-microtubules were polymerized as described for long ones, with the following differences: i) the tubulin concentration in the polymerization solution was increased to 3 µM; ii) the tubulin mixture included 33% (mol/mol) Atto647N-tubulin (for a final fluorescent labeling ratio of 13%); iii) the final volume was 70 µl; iv) polymerization was performed at 37°C for 1 hour; v) resuspension after centrifugation was carried out in BRB80. These short, biotin and brightly Atto647N-labelled microtubules were either used on the same day or, for later use, they were supplemented with 10% glycerol (vol/vol), aliquoted, snap-frozen and stored in liquid-nitrogen. Short microtubules were diluted up to 200-fold in BRB80 before microscopy experiments.

### Microtubule sedimentation assay

1 µM of test protein was mixed with s*tabilized microtubules for microtubule sedimentation assays* (0.5 µM polymerized tubulin) in NuMA microscopy buffer (BRB80, 50 mM KCl, 1 mM MgCl_2_, 1 mM EGTA, 2 mM guanosine-5’-triphosphate [GTP] [Jena Bioscience], 0.15% [wt/vol] methylcellulose, 1% [wt/vol] glucose, 1 mM TCEP). The appropriate amount of KCl was supplemented to achieve a final concentration of 60 mM KCl, accounting for the KCl carried by the various NuMA constructs from their respective storage buffers. The final reaction volume was 30 µl. The mixture was incubated at 30 °C for 15 min. For negative controls, proteins were incubated in the same buffer without microtubules, and microtubules were incubated without proteins. Reaction mixtures were centrifuged in a tabletop centrifuge at 17,000 × *g* for 20 min; 20 µl of supernatant was removed from the top, while the bottom fraction in contact with the pellet was discarded. Pellets were resuspended in 30 µl using NuMA microscopy buffer. Samples were separated by SDS-PAGE and stained using InstantBlue Coomassie protein stain.

### Total Internal Reflection Fluorescence (TIRF) Microscopy

#### Flow chambers for TIRF microscopy assays

##### Chambers with immobilized long or short microtubules

For assays with surface-immobilized GMPCPP-microtubuless, glass coverslips (Menzel coverslips #1.5 18 × 18 mm and 22 × 22 mm, Epredia) were silanized (hexamethyldisilazane [HDMS]) according to a published protocol (Wedler et al., 2022), using HCl (Fisher Scientific) for activation. Chambers were assembled with these silanized, hydrophobic coverslips as described (Gell et al., 2010), using parafilm strips as spacers between two coverslips to create flow channels (18 mm long, ≈3 mm wide, ≈0.1 mm thick).

To immobilize biotinylated GMPCPP-microtubules via NeutrAvidin (Invitrogen), the following sequence of solutions was flowed through the channels at room temperature: i) TetraSpeck microspheres 0.2 μm (Invitrogen) (4.6 × 10^9^ particles ml^-1^ in BRB80, added to allow for channel alignment and drift correction in post-processing of recorded movies), incubated for 2 min; ii) BRB80, twice; iii) 0.4 mg ml^-1^ of NeutrAvidin in BRB80, incubated for 5 min; iv) BRB80; 5% Pluronic F-127 in BRB80, incubated for 10 min; v) BRB80, twice; vi) Atto647N-labeled biotinylated GMPCPP-microtubules, incubated for up to 5 min (depending on the concentration of the stock); vii) assay buffer. For all solutions, the volume was 15 µl, except for microtubules which were suspended in 5 µl. TetraSpeck microspheres and NeutrAvidin dilutions were freshly prepared on the day of the experiment and stored on ice.

##### Chambers with immobilized γTuRC

For γTURC nucleation assays, flow chambers were assembled with biotin-polyethylene glycol-functionalized glass as described previously (Consolati et al., 2022). Channel flowing and γTURC immobilization was performed as described (Brito et al., 2024).

#### TIRF microscopy assays

##### Dynein motility assays

A dynein-dynactin pre-mix of 10 µl was prepared by diluting dynein and dynactin stocks to a molar ratio of 1:2 (dynein complex : dynactin complex) in dynein microscopy buffer (BRB20 [20 mM K-PIPES, 1 mM MgCl_2_, 1 mM EGTA, pH 6.8], supplemented with 1 mM MgCl_2_, 2.5 mM ATP, 0.15% [wt/vol] methylcellulose, 1% glucose [wt/vol], 1 mM TCEP) to a dynein-dynactin concentration approximately six-fold higher than the final concentration. AF546-NuMA^N-term^ and AF647-NuMA^N-term^ stocks were diluted in dynein microscopy buffer; NuMA^FL^, Lis1, and BicD2N^1‒400^ stocks were diluted into BRB20 supplemented by 100 mM KCl and 2 mM TCEP.

For each condition, the appropriate volumes of diluted NuMA, Lis1, and BicD2N^1‒400^ were added to 1.25 or 2.5 µl of dynein-dynactin pre-mix (for Fig. 1 I, J and K, Fig. S2 B [NuMA experiments] or for all the other assays, respectively) to achieve the desired final concentrations of 3‒7 nM dynein, 7-14 nM dynactin, 50‒500 nM AF546-NuMA^N-term^, 50 nM AF647-NuMA^N-term^, 50 nM NuMA^FL^, 10‒5000 nM mCherry-Lis1, 650 nM Lis1 (as stated in the Fig. 1, Fig. S2) in a total volume of 15 µl. The final volume was reached by adding dynein microscopy buffer, supplemented with oxygen scavenger mix (150 µg ml^-1^ catalase, 625 µg ml^-1^ glucose oxidase [SERVA Electrophoresis]) and KCl. The appropriate amount of KCl to reach a final concentration of 40 mM was adjusted for each condition considering the amount of KCl contributed by NuMA, Lis1, and BicD2N^1‒400^ from their respective storage and dilution buffers. For control experiments in the absence of Lis1 or NuMA, their respective dilution buffer were added instead of the protein to maintain the same buffer composition. The final assay mix (15 µl) was flushed all at once in a channel containing immobilized long microtubules at 30 °C in the TIRF microscope incubator (OkoLab).

##### NuMA binding to microtubules elongating from immobilized seeds

NuMA^FL^, NuMA^C-term^ ^L^, NuMA^C-term^ ^S1^, and NuMA^C-term^ ^S2^ pre-mixes were prepared by diluting stock proteins to a concentration 10-fold higher than the final desired concentrations using BRB80, supplemented with 2 mM TCEP and the appropriate amount of KCl to achieve a final concentration of 100 mM KCl, accounting for the KCl carried by the various NuMA constructs from their respective storage buffers. A 150 µM tubulin pre-mix, including 6% (mol/mol) of Atto647N-tubulin (corresponding to a final fluorescent labeling ratio of 2.4%), was prepared in BRB80.

NuMA pre-mixes and the tubulin pre-mix were diluted 10-fold and 15-fold, respectively, into a final volume of 30 µl with NuMA microscopy buffer (see *Microtubule sedimentation assays*), supplemented with oxygen scavenger mix, to reach the final concentrations of: 25-40 nM NuMA^FL^, 10‒75 nM NuMA^C-term^ ^L^, 40‒1000 nM NuMA^C-term^ ^S1^, 25‒75 nM NuMA^C-term^ ^S2^, 10 µM tubulin (as stated in Fig. 2, Fig. 3, Fig. 4, Fig. S4, Fig. S5). For control experiments in the absence of NuMA, NuMA dilution buffer was added instead of the protein to maintain the same buffer composition. The final assay mix (30 µl) was flowed in the microscopy channel in two steps of 15 μl, with a waiting time of 1 min between flows, at 30 °C in the TIRF microscope incubator (for channels with immobilized short stabilized microtubules) or at room temperature (for channels with microtubules elongating from immobilized seeds). For experiments involving “pre-elongated” microtubules (Fig. 4, Fig. S4 C, Fig. S5, Video 1), the initial flush was performed at RT (tubulin only mix), while the second flush (NuMA and tubulin mix) was executed at 30 °C in the TIRF microscope while imaging, after the first ≈10 min of imaging.

##### NuMA binding to microtubules nucleated by immobilized γTuRC

NuMA^FL^ pre-mixes were prepared by diluting the stock protein into BRB80 supplemented with 2 mM TCEP to a concentration 10-fold higher than the final desired concentrations.

NuMA^FL^ pre-mixes were diluted 10-fold into a final volume of 80 µl with γTuRC microscopy buffer (BRB80, 60 mM KCl, 1 mM GTP, 5 mM 2-ME, 0.15% [wt/vol] methylcellulose, 1% [wt/vol] glucose, 0.02% [vol/vol] Brij-35), supplemented with oxygen scavengers (0.1 mg ml^-1^ catalase, 1 mg ml^-1^ glucose oxidase), to reach the final concentrations of 5‒30 nM, as indicated in Fig. 5 and Fig. S6. A tubulin mix, including 38% (mol/mol) of Atto647-tubulin (for a final fluorescent labeling ratio of 5%), was added to the final mix upon addition of NuMA, to reach a final concentration of 10 µM. The final assay mix (60 µl) was flowed in the channel containing immobilized γ-TURC at room temperature, in two steps of 30 μl, with a waiting time of 1 min between flows.

##### NuMA binding to microtubules after laser ablation

NuMA^FL^ and tubulin pre-mixes were diluted and added to the final assay mix as explained in *NuMA binding to immobilized seeds and microtubules elongating from immobilized seeds*, to reach the final concentrations of 40 nM and 10 µM, respectively. The final assay mix (30 µl) was flowed in a channel containing immobilized seeds in two steps of 15 μl, with a waiting time of 1 min between flows, at 30 °C in the TIRF microscope incubator.

After flushing the final assay mix into the channel and starting the imaging, NuMA was allowed to accumulate on the minus end of the GMPCPP-microtubules, while the plus end would start to elongate, for ≈3 min. This facilitated distinguishing the two ends. Subsequently, microtubule laser ablation was performed and, immediately afterwards, a fresh assay mix was flushed once again into the channel while imaging.

##### Microtubule transport by dynein/dynactin/NuMA

Microtubules with NuMA-decorated minus ends in suspension (“lollipop” microtubules) were obtained using the following polymerization mixture: 0.8 µM tubulin, including 33% (mol/mol) of Atto647N-tubulin (for a final fluorescent labeling ratio of 13%); 40 nM NuMA^FL^; 1 mM GMPCPP; 1.4 mM MgCl_2_; 8 mM DTT, BRB80 to reach a final volume of 35 µl. The mixture was incubated 5 min on ice, then 20 min at 37 °C.

Microtubules were centrifuged in a tabletop centrifuge at 17,000 × *g* for 10 min, and the pellet was resuspended in 7 µl of BRB80 supplemented with 10 mM DTT. Lollipops were stored at room temperature and used within an hour.

The dynein-dynactin pre-mix was prepared as described in *Dynein motility assay*; Lis1 pre-mix was prepared by diluting the stock protein 20-fold in BRB20 supplemented by 100 mM KCl and 2 mM TCEP.

Final mix A was prepared as follows: 1.6 µl of Lis1 pre-mix were added to 5 µl of dynein-dynactin pre-mix; the dynein-dynactin-Lis1 concentrated solution was diluted into dynein microscopy buffer to reach the final concentrations of 14 nM dynein, 28 nM dynactin and 1000 nM Lis1 in a final volume of 15 µl. Methylcellulose was omitted from the composition of dynein microscopy buffer to avoid crowding-induced surface localization of lollipop microtubules. The assay buffer was supplemented with oxygen scavenger mix and KCl, adjusted as explained in *Dynein motility assay*. Final mix A (15 µl) was flowed into a channel containing immobilized long microtubules at 30 °C in the TIRF microscope incubator and incubated for up to 5 min to allow accumulation of dynein on microtubules.

During the incubation, final mix B was prepared as described for final mix A, leaving out dynein and introducing up to 4 µl of lollipop microtubules, which were added prior to warming up mix B tube at room temperature.

Assay pre-mixes and final mixes were prepared on ice. BRB80 and BRB20 were prepared as 1× stocks, aliquoted and kept at −20 °C for long-term storage; once defrosted, they were stored at 4 °C for usage up to 2 weeks. Microscopy buffers were freshly prepared for the day of the experiment and stored on ice. After flushing the final assay mix, channels were sealed using vacuum grease (Dow Corning), immediately followed by TIRF microscopy imaging. For the assays which required multiple flushes, the channels were sealed after the last flush.

### TIRF microscopy imaging

For TIRF images shown in Fig. 3 B, C, E (75 nM kymograph) and G, Fig. 4, Fig. 5, Fig. 7, Fig. S4 A, and Video 3, TIRF microscopy was performed using an automated Nikon Eclipse Ti-E with Perfect Focus System, a 100× oil immersion TIRF objective (NA=1.49, CFI SR Apo; Nikon), and Andor iXon 888 Ultra EMCCD cameras (pixel size=100 nm, Andor Technology), controlled by MetaMorph software (Molecular Devices). The sample was excited using 360° TIRF illumination (iLas2, Gataca Systems). The following filter combinations were used: a 405 nm TIRF filter set (TRF49904; Chroma) with an additional ET525/50 (Chroma) bandpass filter; a 488 nm TIRF filter set (TRF49904; Chroma) with an additional ET525/50 (Chroma) bandpass filter; a 561 nm TIRF filter set (TRF49909; Chroma) with additional ET607/70 (Chroma) bandpass filter; a 638 nm TIRF filter set (TRF49909; Chroma) with additional ET607/70 (Chroma) bandpass filter.

For the TIRF images shown in all other figures (except Fig. 1 I and K), Video 1 and Video 2, the imaging was executed on a similar TIRF setup, which differed only for the following components: i) Andor iXon 897 Ultra EMCCD cameras (pixel size=159 nm, Andor Technology); ii) iLas3 laser illumination (Gataca Systems).

For the NuMA-dynein co-localization experiments shown in Fig. 1 I and K, microscopy was carried out using an iMIC (TILL Photonics) TIRF microscope equipped with: a 100× oil immersion objective lens (NA=1.49; Olympus); 1.26× magnification; Evolve 512 EMCCD cameras (original pixel size=160 nm, final pixel size with magnification=127 nm; Photometrics); a quadband filter (405/488/561/640, Semrock). The sample was excited via 360° TIRF illumination, and the system was controlled by Live Analysis software (TILL Photonics).

Typically, snapshots and time-lapse images were acquired with 100 ms exposure for all excited channels (mBFP-γTuRC: 405 nm; mScarlet-NuMA^FL^ and C-terminal constructs: 561 nm; Atto647-tubulin and Atto647N-tubulin: 638 nm), via sequential dual or triple-color imaging. Time-lapses were imaged using an acquisition rate of either: between 3.5 and 6 fps (Fig. 7, Fig. S7, Video 3), 5 fps (Fig. 6, Video 2), 10 fps (Fig. 5), or between 5 and 15 fps (Fig. 3, Fig. 4, Fig. S4, Fig. S5, Video 1). Recording duration was 20‒30 min. For the assays displayed in Fig. 1 C, F, and J and Fig. S2 A, mEGFP-dynein imaging (488 nm excitation) was obtained through stream acquisitions of 2000 frames with 50 ms of exposure. For the NuMA-dynein co-localizations shown in Fig. 1 I and K, visualization was performed by simultaneous dual-color imaging (mEGFP-dynein: 488 nm; mScarlet-NuMA^FL^: 561 nm) acquiring streams of 1000 frames with 100 ms of exposure.

With the exception of the imaging in Fig. 1 I and K, which was done at 18 °C, all other imaging was performed at 30 °C using the OkoLab temperature control.

### TIRF microscopy image processing

All acquired images were processed using Fiji (Schindelin et al., 2012) and FIESTA (Ruhnow, Zwicker and Diez, 2011). Drift correction and channel alignment was performed by tracking the position of TetraSpeck microspheres 0.2 μm using FIESTA.

### Quantifications

#### Dynein processivity

Dynein motility was analyzed by generating kymographs of the mEGFP-dynein signal for all the microtubules in every stream acquisition, using FIESTA. Traces were manually drawn along all detectable straight diagonal lines in a kymograph. For dynein particles that alternated between processive motility and immobile states within the same run without detachment, the entire run was counted as representing an individual processive event. Means and error bars shown in Fig. 1 D and G were obtained via bootstrapping, using a custom MATLAB script. Briefly, events were counted for microtubules selected randomly among all replicates of the same condition, until the total measured microtubule length exceeded 100 µm; the mean number of events/100 µm/100s was computed after 4000 iterations. Dynein velocities were automatically derived by the slope of each processive part in every trace.

##### NuMA’s end selectivity on GMPCPP-microtubules

To assess the tendency of different mScarlet-NuMA constructs to bind the GMPCPP-microtubule end rather than the microtubule lattice in the presence of tubulin (Fig. 2 C and D), the mScarlet-NuMA and Atto647N-tubulin channels were overlaid. Microtubules showing mScarlet signal exclusively at one end were manually counted, their percentage out of all microtubules with any mScarlet signal in one replicate image was calculated, and represented a single data point.

##### Microtubule growth velocity

For each experimental condition shown in Fig. 3, Fig. 4, Fig. S4 and Fig. S5, kymographs were created using FIESTA. The polymerization velocities were obtained by manually drawing a line along the growing ends, and the velocity was automatically calculated from the slope of the line. Minus and plus ends were discerned from one another due to their significant velocity difference, with plus ends always growing faster than minus end. In case the ratio between the velocity of the two ends approximated 1, the microtubule was discarded from the analysis, likely being an antiparallel microtubule pair. When the end velocity was not consistent throughout the movie, particularly in the case of NuMA slowing/stopping the minus end growth only after a certain time, all different velocities for one end were taken into account.

##### NuMA’s binding to γTURC/solution-nucleated microtubules and γTURC-mediated/solution-mediated microtubule nucleation

Binding frequency of NuMA to the minus end of γTURC-nucleated and solution-nucleated microtubules (Fig 5C) was obtained by drawing kymographs of all microtubules in each experiment using FIESTA (line thickness: 8 pixels), and manually counting the number of kymographs where NuMA was found at the microtubule minus end. This included microtubules where NuMA was co-localizing with γTURC already before a detectable microtubule growth. To quantify the maximum mScarlet fluorescence intensity at the minus end of γTURC-nucleated and solution-nucleated microtubules (Fig. 5 D), for each minus end a surrounding area (8×8 pixels) was manually determined. To calculate the maximum mScarlet fluorescence intensity on the surface (Fig. 5 D), areas equal to those drawn around minus ends were identified at randomly chosen sites on the surface (20 per replicate) using a custom macro in Fiji; areas containing microtubules were discarded. For all areas, the maximum mScarlet intensity was then computed and background-corrected using the maximum intensity of an area of equal dimensions on the surface, in the proximity of the minus end/random area of interest (not presenting any mScarlet nor microtubule signal).

The frequency of NuMA adsorption to random microtubule-free areas (8×8 pixels) on the surface (≈20%) was estimated by counting how many times the mScarlet signal was found within the randomly selected surface areas.

For each nucleation assay shown in Fig. S6 A and B, microtubules were manually counted at six different time points using the “Multi-point tool” in Fiji. The total number of γTuRC-nucleated and solution-nucleated microtubules at a given time point was calculated by adding the newly nucleated microtubules to the total from the previous time point. To visualize single-molecule mBFP-γTuRC, the “Z project” function in Fiji was used to generate an average projection of the mBFP channel, which then served as a static background merged with the other channels.

##### NuMA’s binding to laser-ablated GMPCPP-microtubule ends

To assess the propensity of mScarlet-NuMA^FL^ to bind microtubule ends generated by laser ablation (Fig. 6 D), the mScarlet-NuMA and Atto647N-tubulin channels were overlaid. Only the microtubule segments that remained visible on the glass surface after the laser cut (≈37% of total cuts) were included in the analysis. The number of ablation-generated minus and plus ends displaying or lacking mScarlet-NuMA signal was manually counted. The abundance of each type of ablated end was expressed as a percentage of the total number of ablation-generated ends.

##### Dynein-driven lollipop microtubule transport frequency

Landing events were manually counted and the frequency of each type of landing was expressed as a percentage of the total number of lollipops landed throughout the duration of the imaging (Fig 7 E). For moving lollipops, being either anchored to the immobilized microtubule only via their minus end (“single end contact”) or multiple contact points (“aligned parallel, “aligned antiparallel”), a transport event was defined as “processive” if, by drawing a kymograph in the microtubule channel: i) a straight diagonal line associated to the lollipop movement over the immobilized microtubule could be detected, regardless of its length and slope; ii) the line co-localized in the dynein and NuMA channels. The lollipop minus end was identified by the only or most abundant NuMA signal along the lollipop; when identification was not possible, the transport was not counted. The minus end of immobilized microtubules were recognized by the minus-end directed processive motility of dynein and its resulting minus end accumulation. Dynein motility of non-lollipop attached dynein was likely activated by some NuMA that detached from lollipops and became available in solution.

A lollipop was considered as “stuck” if, since its appearance on the immobilized microtubule, its position did not change throughout the whole duration of the movie. Lollipops were labeled as “diffusing” when, upon landing, they wiggled back and forth along the immobilized microtubules, occasionally floating back in solution.

### Data plots

Plots displayed in Fig 1 D and G were obtained with MATLAB, using a standard hyperbolic curve fit. Plots shown in Fig. 1 E and H were produced with the Violin SuperPlot package for MATLAB (Kenny and Schoen, 2021). For all other figures, data plotting was performed using Prism 8 (GraphPad software).

## AUTHOR CONTRIBUTIONS

S.C. and C.M. performed *in vitro* reconstitution experiments and analyzed data. S.C. generated the data shown in the figures and designed the figures. S.C., C.M. and S.S. performed protein purifications. F.R. facilitated and performed data analysis and provided microscopy support. S.C., C.M., S.S. and M.G. cloned protein expression constructs and expressed proteins in insect cells or bacteria. C.B. provided reagents and support for experiments with γTuRC. T.S. supervised and coordinated the project. S.C. and T.S. wrote the paper.

## ACKNOWLEDGEMENTS

We thank Marina Mapelli for the gift of a codon-optimized full-length NuMA plasmid, Julian Gannon and Raquel Garcia-Castellanos for cloning, protein expression and purification support, Davide Normanno for microscopy support, the UAB (Universitat Autonoma Barcelona) Microscopy and X-ray Diffraction Service for negative stain electron microscopy. We acknowledge support of the Spanish Ministry of Science and Innovation through the Centro de Excelencia Severo Ochoa (CEX2020-001049-S, MCIN/AEI /10.13039/501100011033), and the Generalitat de Catalunya through the CERCA programme. We are grateful to the CRG Core Technologies Units for their support and assistance.

## REFERENCES

Aher, A., Urnavicius, L., Xue, A., Neselu, K. and Kapoor, T.M. (2024) ‘Structure of the γ-tubulin ring complex-capped microtubule’, Nature Structural & Molecular Biology, 31(7), pp. 1124–1133. Available at: 10.1038/s41594-024-01264-z.

Brito, C., Serna, M., Guerra, P., Llorca, O. and Surrey, T. (2024) ‘Transition of human γ-tubulin ring complex into a closed conformation during microtubule nucleation’, Science, 383(6685), pp. 870–876. Available at: 10.1126/science.adk6160.

Canty, J.T., Hensley, A., Aslan, M., Jack, A. and Yildiz, A. (2023) ‘TRAK adaptors regulate the recruitment and activation of dynein and kinesin in mitochondrial transport’, Nature Communications, 14(1). Available at: 10.1038/S41467-023-36945-8.

Canty, J.T., Tan, R., Kusakci, E., Fernandes, J. and Yildiz, A. (2021) ‘Structure and Mechanics of Dynein Motors’, Annual Review of Biophysics, 50(1), pp. 549–574. Available at: 10.1146/annurev-biophys-111020-101511.

Canty, J.T. and Yildiz, A. (2020) ‘Activation and Regulation of Cytoplasmic Dynein’, Trends in Biochemical Sciences, 45(5), pp. 440–453. Available at: 10.1016/j.tibs.2020.02.002.

Carter, A.P., Diamant, A.G. and Urnavicius, L. (2016) ‘How dynein and dynactin transport cargos: a structural perspective’. Available at: 10.1016/j.sbi.2015.12.003.

Castoldi, M. and Popov, A. V. (2003) ‘Purification of brain tubulin through two cycles of polymerization-depolymerization in a high-molarity buffer’, Protein Expression and Purification, 32(1), pp. 83–88. Available at: 10.1016/S1046-5928(03)00218-3.

Chaaban, S. and Carter, A.P. (2022) ‘Structure of dynein-dynactin on microtubules shows tandem adaptor binding’, Nature, 610(7930), pp. 212–216. Available at: 10.1038/S41586-022-05186-Y.

Chang, C.-C., Huang, T.-L., Shimamoto, Y., Tsai, S.-Y. and Hsia, K.-C. (2017) ‘Regulation of mitotic spindle assembly factor NuMA by Importin-β’, Journal of Cell Biology, 216(11), pp. 3453–3462. Available at: 10.1083/jcb.201705168.

Chowdhury, S., Ketcham, S.A., Schroer, T.A. and Lander, G.C. (2015) ‘Structural organization of the dynein-dynactin complex bound to microtubules’, Nature structural & molecular biology, 22(4), pp. 345–347. Available at: 10.1038/NSMB.2996.

Compton, D.A. and Luo, C. (1995) ‘Mutation of the predicted p34cdc2 phosphorylation sites in numa impair the assembly of the mitotic spindle and block mitosis’, Journal of Cell Science, 108(2), pp. 621–633. Available at: 10.1242/jcs.108.2.621.

Consolati, T., Henkin, G., Roostalu, J. and Surrey, T. (2022) ‘Real-Time Imaging of Single γTuRC-Mediated Microtubule Nucleation Events In Vitro by TIRF Microscopy’, Methods in Molecular Biology, 2430, pp. 315–336. Available at: 10.1007/978-1-0716-1983-4_21.

Consolati, T., Locke, J., Roostalu, J., Chen, Z.A., Gannon, J., Asthana, J., Lim, W.M., Martino, F., Cvetkovic, M.A., Rappsilber, J., Costa, A. and Surrey, T. (2020) ‘Microtubule Nucleation Properties of Single Human γTuRCs Explained by Their Cryo-EM Structure’, Developmental Cell, 53(5), pp. 603–617.e8. Available at: 10.1016/j.devcel.2020.04.019.

D’amico, E.A., Ahmad, M.U.D., Cmentowski, V., Girbig, M., Müller, F., Wohlgemuth, S., Brockmeyer, A., Maffini, S., Janning, P., Vetter, I.R., Carter, A.P., Perrakis, A. and Musacchio, A. (2022) ‘Conformational transitions of the Spindly adaptor underlie its interaction with Dynein and Dynactin’, Journal of Cell Biology, 221(11). Available at: 10.1083/jcb.202206131.

Dendooven, T., Yatskevich, S., Burt, A., Chen, Z.A., Bellini, D., Rappsilber, J., Kilmartin, J. V. and Barford, D. (2024) ‘Structure of the native γ-tubulin ring complex capping spindle microtubules’, Nature Structural & Molecular Biology, 31(7), pp. 1134–1144. Available at: 10.1038/s41594-024-01281-y.

Drechsler, H. and McAinsh, A.D. (2016) ‘Kinesin-12 motors cooperate to suppress microtubule catastrophes and drive the formation of parallel microtubule bundles’, Proceedings of the National Academy of Sciences, 113(12), pp. E1635–E1644. Available at: 10.1073/pnas.1516370113.

Du, Q., Taylor, L., Compton, D.A. and Macara, I.G. (2002) ‘LGN Blocks the Ability of NuMA to Bind and Stabilize Microtubules’, Current Biology, 12(22), pp. 1928–1933. Available at: 10.1016/S0960-9822(02)01298-8.

Elshenawy, M.M., Kusakci, E., Volz, S., Baumbach, J., Bullock, S.L. and Yildiz, A. (2020) ‘Lis1 activates dynein motility by modulating its pairing with dynactin’, Nature cell biology, 22(5), p. 570. Available at: 10.1038/S41556-020-0501-4.

Fink, G., Hajdo, L., Skowronek, K.J., Reuther, C., Kasprzak, A.A. and Diez, S. (2009) ‘The mitotic kinesin-14 Ncd drives directional microtubule–microtubule sliding’, Nature Cell Biology, 11(6), pp. 717–723. Available at: 10.1038/ncb1877.

Forth, S., Hsia, K.C., Shimamoto, Y. and Kapoor, T.M. (2014) ‘Asymmetric friction of nonmotor MAPs can lead to their directional motion in active microtubule networks’, Cell, 157(2), pp. 420–432. Available at: 10.1016/j.cell.2014.02.018.

Gaglio, T., Saredi, A. and Compton, D.A. (1995) ‘NuMA is required for the organization of microtubules into aster-like mitotic arrays.’, Journal of Cell Biology, 131(3), pp. 693–708. Available at: 10.1083/JCB.131.3.693.

Gallini, S., Carminati, M., De Mattia, F., Pirovano, L., Martini, E., Oldani, A., Asteriti, I.A., Guarguaglini, G. and Mapelli, M. (2016) ‘NuMA phosphorylation by aurora-a orchestrates spindle orientation’, Current Biology, 26(4), pp. 458–469. Available at: 10.1016/j.cub.2015.12.051.

Gama, J.B., Pereira, C., Simões, P.A., Celestino, R., Reis, R.M., Barbosa, D.J., Pires, H.R., Carvalho, C., Amorim, J., Carvalho, A.X., Cheerambathur, D.K. and Gassmann, R. (2017) ‘Molecular mechanism of dynein recruitment to kinetochores by the Rod-Zw10-Zwilch complex and Spindly’, Journal of Cell Biology, 216(4), pp. 943–960. Available at: 10.1083/JCB.201610108/VIDEO-4.

Gell, C., Bormuth, V., Brouhard, G.J., Cohen, D.N., Diez, S., Friel, C.T., Helenius, J., Nitzsche, B., Petzold, H., Ribbe, J., Schäffer, E., Stear, J.H., Trushko, A., Varga, V., Widlund, P.O., Zanic, M. and Howard, J. (2010) ‘Microtubule dynamics reconstituted in vitro and imaged by single-molecule fluorescence microscopy’, Methods in cell biology, 95(C), pp. 221–245. Available at: 10.1016/S0091-679X(10)95013-9.

Harborth, J., Wang, J., Gueth-Hallonet, C., Weber, K. and Osborn, M. (1999) ‘Self assembly of NuMA: Multiarm oligomers as structural units of a nuclear lattice’, EMBO Journal, 18(6), pp. 1689–1700. Available at: 10.1093/emboj/18.6.1689.

Harborth, J., Weber, K. and Osborn, M. (1995) ‘Epitope mapping and direct visualization of the parallel, in-register arrangement of the double-stranded coiled-coil in the NuMA protein.’, The EMBO Journal, 14(11), pp. 2447–2460. Available at: 10.1002/j.1460-2075.1995.tb07242.x.

Haren, L. and Merdes, A. (2002) ‘Direct binding of NuMA to tubulin is mediated by a novel sequence motif in the tail domain that bundles and stabilizes microtubules’, Journal of Cell Science, 115(9), pp. 1815–1824. Available at: 10.1242/jcs.115.9.1815.

Heald, R., Tournebize, R., Blank, T., Sandaltzopoulos, R., Becker, P., Hyman, A. and Karsenti, E. (1996) ‘Self-organization of microtubules into bipolar spindles around artificial chromosomes in Xenopus egg extracts’, Nature, 382(6590), pp. 420–425. Available at: 10.1038/382420a0.

Heald, R., Tournebize, R., Habermann, A., Karsenti, E. and Hyman, A. (1997) ‘Spindle Assembly in Xenopus Egg Extracts: Respective Roles of Centrosomes and Microtubule Self-Organization’, 138(3), pp. 615–628.

Henkin, G., Chew, W.-X., Nédélec, F. and Surrey, T. (2022) ‘Cross-linker design determines microtubule network organization by opposing motors’, Proceedings of the National Academy of Sciences, 119(33). Available at: 10.1073/pnas.2206398119.

Hentrich, C. and Surrey, T. (2010) ‘Microtubule organization by the antagonistic mitotic motors kinesin-5 and kinesin-14’, The Journal of Cell Biology, 189(3), pp. 465–480. Available at: 10.1083/jcb.200910125.

Hong, W., Takshak, A., Osunbayo, O., Kunwar, A. and Vershinin, M. (2016) ‘The Effect of Temperature on Microtubule-Based Transport by Cytoplasmic Dynein and Kinesin-1 Motors’, Biophysical Journal, 111(8), p. 1816. Available at: 10.1016/j.bpj.2016.09.040.

Htet, Z.M., Gillies, J.P., Baker, R.W., Leschziner, A.E., DeSantis, M.E. and Reck-Peterson, S.L. (2020) ‘Lis1 promotes the formation of activated cytoplasmic dynein-1 complexes’, Nature cell biology, 22(5), p. 518. Available at: 10.1038/S41556-020-0506-Z.

Hueschen, C.L., Galstyan, V., Amouzgar, M., Phillips, R. and Dumont, S. (2019) ‘Microtubule End-Clustering Maintains a Steady-State Spindle Shape’, Current Biology, 29(4), pp. 700–708.e5. Available at: 10.1016/j.cub.2019.01.016.

Hueschen, C.L., Kenny, S.J., Xu, K. and Dumont, S. (2017) ‘NuMA recruits dynein activity to microtubule minus-ends at mitosis’, eLife, 6(e29328), pp. 1–26. Available at: 10.7554/eLife.29328.

Hyman, A.A. (1991) ‘Preparation of marked microtubules for the assay of the polarity of microtubule-based motors by fluorescence’, in Journal of Cell Science. J Cell Sci Suppl, pp. 125–127. Available at: 10.1242/jcs.1991.supplement_14.25.

Jha, R., Roostalu, J., Cade, N.I., Trokter, M. and Surrey, T. (2017) ‘Combinatorial regulation of the balance between dynein microtubule end accumulation and initiation of directed motility’, The EMBO Journal, 36(22), pp. 3387– 3404. Available at: 10.15252/embj.201797077.

Jiang, K., Rezabkova, L., Hua, S., Liu, Q., Capitani, G., Altelaar, A.F.M., Heck, A.J.R., Kammerer, R.A., Steinmetz, M.O. and Akhmanova, A. (2017) ‘Microtubule minus-end regulation at spindle poles by an ASPM-katanin complex’, Nature Cell Biology, 19(5), pp. 480–492. Available at: 10.1038/ncb3511.

Jin, M., Pomp, O., Shinoda, T., Toba, S., Torisawa, T., Furuta, K., Oiwa, K., Yasunaga, T., Kitagawa, D., Matsumura, S., Miyata, T., Tan, T.T., Reversade, B. and Hirotsune, S. (2017) ‘Katanin p80, NuMA and cytoplasmic dynein cooperate to control microtubule dynamics’, Scientific reports, 7. Available at: 10.1038/SREP39902.

Kapitein, L.C., Peterman, E.J.G., Kwok, B.H., Kim, J.H., Kapoor, T.M. and Schmidt, C.F. (2005) ‘The bipolar mitotic kinesin Eg5 moves on both microtubules that it crosslinks’, Nature, 435(7038), pp. 114–118. Available at: 10.1038/nature03503.

Karasmanis, E.P., Reimer, J.M., Kendrick, A.A., Nguyen, K.H.V., Rodriguez, J.A., Truong, J.B., Lahiri, I., Reck-Peterson, S.L. and Leschziner, A.E. (2023) ‘Lis1 relieves cytoplasmic dynein-1 autoinhibition by acting as a molecular wedge’, Nature Structural and Molecular Biology, 30(9), pp. 1357–1364. Available at: 10.1038/s41594-023-01069-6.

Kashina, A.S., Baskin, R.J., Cole, D.G., Wedaman, K.P., Saxton, W.M. and Scholey, J.M. (1996) ‘A bipolar kinesin’, Nature, 379(6562), pp. 270–272. Available at: 10.1038/379270a0.

Kendrick, A.A., Dickey, A.M., Redwine, W.B., Tran, P.T., Vaites, L.P., Dzieciatkowska, M., Harper, J.W. and Reck-Peterson, S.L. (2019) ‘Hook3 is a scaffold for the opposite-polarity microtubule-based motors cytoplasmic dynein-1 and KIF1C’, Journal of Cell Biology, 218(9), pp. 2982–3001. Available at: 10.1083/JCB.201812170.

Kenny, M. and Schoen, I. (2021) ‘Violin SuperPlots: visualizing replicate heterogeneity in large data sets’, Molecular Biology of the Cell, 32(15), p. 1333. Available at: 10.1091/MBC.E21-03-0130.

Kettenbach, A.N., Schweppe, D.K., Faherty, B.K., Pechenick, D., Pletnev, A.A. and Gerber, S.A. (2011) ‘Quantitative Phosphoproteomics Identifies Substrates and Functional Modules of Aurora and Polo-Like Kinase Activities in Mitotic Cells’, Science Signaling, 4(179). Available at: 10.1126/scisignal.2001497.

Kotak, S., Afshar, K., Busso, C. and Gönczy, P. (2016) ‘Aurora A kinase regulates proper spindle positioning in C. elegans and in human cells’, Journal of cell science, 129(15), pp. 3015–3025. Available at: 10.1242/JCS.184416.

Kotak, S., Busso, C. and Gönczy, P. (2012) ‘Cortical dynein is critical for proper spindle positioning in human cells’, Journal of Cell Biology, 199(1), pp. 97–110. Available at: 10.1083/jcb.201203166.

Kotak, S., Busso, C. and Gönczy, P. (2013) ‘NuMA phosphorylation by CDK1 couples mitotic progression with cortical dynein function’, The EMBO Journal, 32(18), pp. 2517–2529. Available at: 10.1038/emboj.2013.172.

Lee, I.G., Cason, S.E., Alqassim, S.S., Holzbaur, E.L.F. and Dominguez, R. (2020) ‘A tunable LIC1-adaptor interaction modulates dynein activity in a cargo-specific manner’, Nature communications, 11(1). Available at: 10.1038/S41467-020-19538-7.

Liu, Y., Salter, H.K., Holding, A.N., Johnson, C.M., Stephens, E., Lukavsky, P.J., Walshaw, J. and Bullock, S.L. (2013) ‘Bicaudal-D uses a parallel, homodimeric coiled coil with heterotypic registry to coordinate recruitment of cargos to dynein’, Genes & Development, 27(11), p. 1233. Available at: 10.1101/GAD.212381.112.

Lydersen, B.K. and Pettijohn, D.E. (1980) ‘Human-specific nuclear protein that associates with the polar region of the mitotic apparatus: Distribution in a human/hamster hybrid cell’, Cell, 22(2), pp. 489–499. Available at: 10.1016/0092-8674(80)90359-1.

Maekawa, T., Leslie, R. and Kuriyama, R. (1991) ‘Identification of a minus end-specific microtubule-associated protein located at the mitotic poles in cultured mammalian cells’, European Journal of Cell Biology, 54(2), pp. 255–267. Available at: https://experts.umn.edu/en/publications/identification-of-a-minus-end-specific-microtubule-associated-pro (Accessed: 29 July 2024).

Marzo, M.G., Griswold, J.M. and Markus, S.M. (2020) ‘Pac1/LIS1 stabilizes an uninhibited conformation of dynein to coordinate its localization and activity’, Nature cell biology, 22(5), p. 559. Available at: 10.1038/S41556-020-0492-1.

McKenney, R.J., Huynh, W., Tanenbaum, M.E., Bhabha, G. and Vale, R.D. (2014) ‘Activation of cytoplasmic dynein motility by dynactin-cargo adapter complexes’, Science, 345(6194), pp. 337–341. Available at: 10.1126/science.1254198.

McNally, F.J. and Roll-Mecak, A. (2018) ‘Microtubule-severing enzymes: From cellular functions to molecular mechanism’, Journal of Cell Biology, 217(12), pp. 4057–4069. Available at: 10.1083/jcb.201612104.

Merdes, A., Heald, R., Samejima, K., Earnshaw, W.C. and Cleveland, D.W. (2000) ‘Formation of spindle poles by dynein/dynactin-dependent transport of NuMA’, Journal of Cell Biology, 149(4), pp. 851–861. Available at: 10.1083/jcb.149.4.851.

Merdes, A., Ramyar, K., Vechio, J.D. and Cleveland, D.W. (1996) ‘A complex of NuMA and cytoplasmic dynein is essential for mitotic spindle assembly’, Cell, 87(3), pp. 447–458. Available at: 10.1016/S0092-8674(00)81365-3.

Monda, J.K. and Cheeseman, I.M. (2018) ‘Nde1 promotes diverse dynein functions through differential interactions and exhibits an isoform-specific proteasome association’, Molecular Biology of the Cell. Edited by K.G. Kozminski, 29(19), pp. 2336–2345. Available at: 10.1091/mbc.E18-07-0418.

Moritz, M., Braunfeld, M.B., Guénebaut, V., Heuser, J. and Agard, D.A. (2000) ‘Structure of the γ-tubulin ring complex: a template for microtubule nucleation’, Nature Cell Biology, 2(6), pp. 365–370. Available at: 10.1038/35014058.

Nachury, M. V., Maresca, T.J., Salmon, W.C., Waterman-Storer, C.M., Heald, R. and Weis, K. (2001) ‘Importin β is a mitotic target of the small GTPase ran in spindle assembly’, Cell, 104(1), pp. 95–106. Available at: 10.1016/S0092-8674(01)00194-5.

Okumura, M., Natsume, T., Kanemaki, M.T. and Kiyomitsu, T. (2018) ‘Dynein–dynactin–NuMA clusters generate cortical spindle-pulling forces as a multiarm ensemble’, eLife, 7, pp. 1–24. Available at: 10.7554/eLife.36559.

Olenick, M.A. and Holzbaur, E.L.F. (2019) ‘Dynein activators and adaptors at a glance’, Journal of Cell Science, 132(6), pp. 1–7. Available at: 10.1242/jcs.227132.

Petry, S., Groen, A.C., Ishihara, K., Mitchison, T.J. and Vale, R.D. (2013) ‘Branching Microtubule Nucleation in Xenopus Egg Extracts Mediated by Augmin and TPX2’, Cell, 152(4), pp. 768–777. Available at: 10.1016/j.cell.2012.12.044.

Pirovano, L., Culurgioni, S., Carminati, M., Alfieri, A., Monzani, S., Cecatiello, V., Gaddoni, C., Rizzelli, F., Foadi, J., Pasqualato, S. and Mapelli, M. (2019) ‘Hexameric NuMA:LGN structures promote multivalent interactions required for planar epithelial divisions’, Nature Communications, 10(1). Available at: 10.1038/s41467-019-09999-w.

Qiu, R., Zhang, J. and Xiang, X. (2019) ‘LIS1 regulates cargo-adapter–mediated activation of dynein by overcoming its autoinhibition in vivo’, Journal of Cell Biology, 218(11), pp. 3630–3646. Available at: 10.1083/JCB.201905178.

Raaijmakers, J.A. and Medema, R.H. (2014) ‘Function and regulation of dynein in mitotic chromosome segregation’, Chromosoma, 123(5), pp. 407–422. Available at: 10.1007/s00412-014-0468-7.

Rai, D., Song, Y., Hua, S., Stecker, K., Monster, J.L., Yin, V., Stucchi, R., Xu, Y., Zhang, Y., Chen, F., Katrukha, E.A., Altelaar, M., Heck, A.J.R., Wieczorek, M., Jiang, K. and Akhmanova, A. (2024) ‘CAMSAPs and nucleation-promoting factors control microtubule release from γ-TuRC’, Nature Cell Biology, 26(3), p. 404. Available at: 10.1038/S41556-024-01366-2.

Reck-Peterson, S.L., Redwine, W.B., Vale, R.D. and Carter, A.P. (2018) ‘The cytoplasmic dynein transport machinery and its many cargoes’, Nature Reviews Molecular Cell Biology, 19(6), pp. 382–398. Available at: 10.1038/s41580-018-0004-3.

Redwine, W.B., DeSantis, M.E., Hollyer, I., Htet, Z.M., Tran, P.T., Swanson, S.K., Florens, L., Washburn, M.P. and Reck-Peterson, S.L. (2017) ‘The human cytoplasmic dynein interactome reveals novel activators of motility’, eLife, 6. Available at: 10.7554/ELIFE.28257.

Renna, C., Rizzelli, F., Carminati, M., Gaddoni, C., Pirovano, L., Cecatiello, V., Pasqualato, S. and Mapelli, M. (2020) ‘Organizational Principles of the NuMA-Dynein Interaction Interface and Implications for Mitotic Spindle Functions’, Structure, 28(7), pp. 820–829.e6. Available at: 10.1016/j.str.2020.04.017.

Ruhnow, F., Kloß, L. and Diez, S. (2017) ‘Challenges in Estimating the Motility Parameters of Single Processive Motor Proteins’, Biophysical Journal, 113(11), pp. 2433–2443. Available at: 10.1016/j.bpj.2017.09.024.

Ruhnow, F., Zwicker, D. and Diez, S. (2011) ‘Tracking Single Particles and Elongated Filaments with Nanometer Precision’, Biophysical Journal, 100(11), p. 2820. Available at: 10.1016/J.BPJ.2011.04.023.

Sana, S., Keshri, R., Rajeevan, A., Kapoor, S. and Kotak, S. (2018) ‘Plk1 regulates spindle orientation by phosphorylating NuMA in human cells’, Life Science Alliance, 1(6), p. e201800223. Available at: 10.26508/lsa.201800223.

Schindelin, J., Arganda-Carreras, I., Frise, E., Kaynig, V., Longair, M., Pietzsch, T., Preibisch, S., Rueden, C., Saalfeld, S., Schmid, B., Tinevez, J.Y., White, D.J., Hartenstein, V., Eliceiri, K., Tomancak, P. and Cardona, A. (2012) ‘Fiji: an open-source platform for biological-image analysis’, Nature methods, 9(7), pp. 676–682. Available at: 10.1038/NMETH.2019.

Schlager, M.A., Hoang, H.T., Urnavicius, L., Bullock, S.L. and Carter, A.P. (2014) ‘ In vitro reconstitution of a highly processive recombinant human dynein complex’, The EMBO Journal, 33(17), pp. 1855–1868. Available at: 10.15252/embj.201488792.

Schroeder, C.M. and Vale, R.D. (2016) ‘Assembly and activation of dynein–dynactin by the cargo adaptor protein Hook3’, Journal of Cell Biology, 214(3), pp. 309–318. Available at: 10.1083/JCB.201604002.

Seldin, L., Poulson, N.D., Foote, H.P. and Lechler, T. (2013) ‘NuMA localization, stability, and function in spindle orientation involve 4.1 and Cdk1 interactions’, Molecular Biology of the Cell. Edited by E. Holzbaur, 24(23), pp. 3651–3662. Available at: 10.1091/mbc.e13-05-0277.

Sikirzhytski, V., Magidson, V., Steinman, J.B., He, J., Le Berre, M., Tikhonenko, I., Ault, J.G., McEwen, B.F., Chen, J.K., Sui, H., Piel, M., Kapoor, T.M. and Khodjakov, A. (2014) ‘Direct kinetochore–spindle pole connections are not required for chromosome segregation’, Journal of Cell Biology, 206(2), pp. 231–243. Available at: 10.1083/jcb.201401090.

Silk, A.D., Holland, A.J. and Cleveland, D.W. (2009) ‘Requirements for NuMA in maintenance and establishment of mammalian spindle poles’, Journal of Cell Biology, 184(5), pp. 677–690. Available at: 10.1083/jcb.200810091.

Singh, K., Lau, C.K., Manigrasso, G., Gama, J.B., Gassmann, R. and Carter, A.P. (2023) ‘Molecular mechanism of dynein-dynactin activation by JIP3 and LIS1’, bioRxiv, p. 2022.08.17.504273. Available at: 10.1101/2022.08.17.504273.

Snapp, E.L., Hegde, R.S., Francolini, M., Lombardo, F., Colombo, S., Pedrazzini, E., Borgese, N. and Lippincott-Schwartz, J. (2003) ‘Formation of stacked ER cisternae by low affinity protein interactions’, Journal of Cell Biology, 163(2), pp. 257–269. Available at: 10.1083/JCB.200306020.

So, C., Menelaou, K., Uraji, J., Harasimov, K., Steyer, A.M., Seres, K.B., Bucevičius, J., Lukinavičius, G., Möbius, W., Sibold, C., Tandler-Schneider, A., Eckel, H., Moltrecht, R., Blayney, M., Elder, K. and Schuh, M. (2022) ‘Mechanism of spindle pole organization and instability in human oocytes’, Science, 375(6581). Available at: 10.1126/science.abj3944.

Sun, M., Jia, M., Ren, H., Yang, B., Chi, W., Xin, G., Jiang, Q. and Zhang, C. (2021) ‘NuMA regulates mitotic spindle assembly, structural dynamics and function via phase separation’, Nature Communications, 12(1). Available at: 10.1038/s41467-021-27528-6.

Toorn, M. van, Gooch, A., Boerner, S. and Kiyomitsu, T. (2022) ‘NuMA Deficiency Causes Micronuclei via Checkpoint-Insensitive k-Fiber Minus-End Detachment From Mitotic Spindle Poles’, SSRN Electronic Journal, pp. 1–9. Available at: 10.2139/ssrn.4233516.

Torisawa, T., Ichikawa, M., Furuta, A., Saito, K., Oiwa, K., Kojima, H., Toyoshima, Y.Y. and Furuta, K. (2014) ‘Autoinhibition and cooperative activation mechanisms of cytoplasmic dynein’, Nature cell biology, 16(11), pp. 1118–1124. Available at: 10.1038/NCB3048.

Tsuchiya, K., Hayashi, H., Nishina, M., Okumura, M., Sato, Y., Kanemaki, M.T., Goshima, G. and Kiyomitsu, T. (2020) ‘Ran-GTP Is Non-essential to Activate NuMA for Mitotic Spindle-Pole Focusing but Dynamically Polarizes HURP Near Chromosomes’, Current Biology, 31(1), pp. 115–127.e3. Available at: 10.1016/j.cub.2020.09.091.

Uehara, R., Nozawa, R.S., Tomioka, A., Petry, S., Vale, R.D., Obuse, C. and Goshima, G. (2009) ‘The augmin complex plays a critical role in spindle microtubule generation for mitotic progression and cytokinesis in human cells’, Proceedings of the National Academy of Sciences of the United States of America, 106(17), pp. 6998–7003. Available at: 10.1073/PNAS.0901587106/SUPPL_FILE/0901587106SI.PDF.

Urnavicius, L., Lau, C.K., Elshenawy, M.M., Morales-Rios, E., Motz, C., Yildiz, A. and Carter, A.P. (2018) ‘Cryo-EM shows how dynactin recruits two dyneins for faster movement’, Nature, 554(7691), pp. 202–206. Available at: 10.1038/nature25462.

Urnavicius, L., Zhang, K., Diamant, A.G., Motz, C., Schlager, M.A., Yu, M., Patel, N.A., Robinson, C. V. and Carter, A.P. (2015) ‘The structure of the dynactin complex and its interaction with dynein’, Science, 347(6229), pp. 1441–1446. Available at: 10.1126/science.aaa4080.

Verde, F., Berrez, J.-M., Antony, C. and Karsenti, E. (1991) ‘Taxol-induced Microtubule Asters in Mitotic Extracts of Xenopus Eggs’:, Journal of Cell Biology, 112(6), pp. 1177–1187.

Wang, S., Ketcham, S.A., Schön, A., Goodman, B., Wang, Y., Yates, J., Freire, E., Schroer, T.A. and Zheng, Y. (2013) ‘Nudel/NudE and Lis1 promote dynein and dynactin interaction in the context of spindle morphogenesis’, Molecular Biology of the Cell. Edited by W. Bement, 24(22), pp. 3522–3533. Available at: 10.1091/mbc.e13-05-0283.

Wasilko, D.J., Edward Lee, S., Stutzman-Engwall, K.J., Reitz, B.A., Emmons, T.L., Mathis, K.J., Bienkowski, M.J., Tomasselli, A.G. and David Fischer, H. (2009) ‘The titerless infected-cells preservation and scale-up (TIPS) method for large-scale production of NO-sensitive human soluble guanylate cyclase (sGC) from insect cells infected with recombinant baculovirus’, Protein Expression and Purification, 65(2), pp. 122–132. Available at: 10.1016/J.PEP.2009.01.002.

Wedler, V., Quinones, D., Peisert, H. and Schäffer, E. (2022) ‘A Quick and Reproducible Silanization Method by Using Plasma Activation for Hydrophobicity-Based Kinesin Single Molecule Fluorescence–Microscopy Assays’, Chemistry – A European Journal, 28(64), p. e202202036. Available at: 10.1002/CHEM.202202036.

Weissmann, F., Petzold, G., VanderLinden, R., Huis in’t Veld, P.J., Brown, N.G., Lampert, F., Westermann, S., Stark, H., Schulman, B.A. and Peters, J.-M. (2016) ‘biGBac enables rapid gene assembly for the expression of large multisubunit protein complexes’, Proceedings of the National Academy of Sciences, 113(19), pp. E2564–E2569. Available at: 10.1073/pnas.1604935113.

Wieczorek, M., Ti, S.C., Urnavicius, L., Molloy, K.R., Aher, A., Chait, B.T. and Kapoor, T.M. (2021) ‘Biochemical reconstitutions reveal principles of human γ-TuRC assembly and function’, Journal of Cell Biology, 220(3). Available at: 10.1083/JCB.202009146/211719.

Wiese, C., Wilde, A., Moore, M.S., Adam, S.A., Merdes, A. and Zheng, Y. (2001) ‘Role of Importin-β in Coupling Ran to Downstream Targets in Microtubule Assembly’, Science, 291(5504), pp. 653–656. Available at: 10.1126/science.1057661.

Yildiz, A. and Zhao, Y. (2023) ‘Dyneins’, Current Biology, 33(24), pp. R1274–R1279. Available at: 10.1016/j.cub.2023.10.064.

Zacharias, D.A., Violin, J.D., Newton, A.C. and Tsien, R.Y. (2002) ‘Partitioning of Lipid-Modified Monomeric GFPs into Membrane Microdomains of Live Cells’, Science, 296(5569), pp. 913–916. Available at: 10.1126/science.1068539.

Zhang, K., Foster, H.E., Rondelet, A., Lacey, S.E., Bahi-Buisson, N., Bird, A.W. and Carter, A.P. (2017) ‘Cryo-EM Reveals How Human Cytoplasmic Dynein Is Auto-inhibited and Activated’, Cell, 169(7), pp. 1303–1314.e18. Available at: 10.1016/j.cell.2017.05.025.

Zhao, Y., Oten, S. and Yildiz, A. (2023) ‘Nde1 promotes Lis1-mediated activation of dynein’, Nature Communications, 14(1), pp. 1–14. Available at: 10.1038/s41467-023-42907-x.

Zheng, Y., Wong, M.L., Alberts, B. and Mitchison, T. (1995) ‘Nucleation of microtubule assembly by a gamma-tubulin-containing ring complex’, Nature, 378(6557), pp. 578–583. Available at: 10.1038/378578A0.

