## Supplementary Figures and Video legends for "NuMA is a mitotic adaptor protein that activates dynein and connects it to microtubule minus ends"

#### SUPPLEMENTARY FIGURE LEGENDS

**Fig. S1 | Purified proteins used in dynein motility assays. (A)** Coomassie Blue-stained SDS-PAGE of the purified proteins. The expected molecular weight (kDa) of each purified protein is indicated. For dynein and dynactin complexes, the size and name of each subunit is specified. Western blots (dashed frames) were performed to detect the smallest dynein subunits (not visible on SDS-PAGE) and verify the identity of some dynactin SDS-PAGE bands. **(B)** Analysis of the oligomeric state of each purified protein by mass photometry. The molecular weight, with the associated standard deviation (σ), of the most abundant species present in each sample is indicated in the mass histograms. The symmetric parts of the histograms around mass 0 kDa represent the background signal of the buffer. For AF647-NuMA$^{\text{N-term}}$, the 162 kDa peak may represent the oligomerized state of a minor contaminant of 60–65 kDa. **C.** Representative negative staining electron microscopy images of mScarlet-NuMA$^{\text{FL}}$ particles. NuMA appears as an elongated, thin rod; the size corresponds to the expected length of its coiled coil region (207 ± 20 nm (Harborth, Weber and Osborn, 1995)). Yellow arrowheads indicate the extremities of the rod. At one end, two globular heads are visible (N-terminal Hook domains). The absence of significant bending suggests lack of auto-inhibition. Spherical particles adsorbed on the surface are likely micelles of Brij-35 (≈9 nm), which is present above its CMC (critical micelle concentration) in the storage buffer.

**Fig. S2 | Effect of Lis1 and temperature on dynein motility in the presence of adaptors. (A)** Representative kymographs showing the motility of mEGFP-dynein on immobilized GMPCPP-microtubules in the presence of dynactin and BicD2N$^{1\text{–}400}$ at different mCherry-Lis1 concentrations. Lis1 stimulates BicD2N$^{1\text{–}400}$-induced processive motility in a dose-dependent manner. **(B)** Velocity distributions of mEGFP-dynein processive runs (mean ± SD) in the presence of either mScarlet-NuMA$^{\text{N-term}}$, mScarlet-NuMA$^{\text{FL}}$ or BicD2N$^{1\text{–}400}$, as indicated. Each circle corresponds to a single velocity; $n$ = 49, 55, 50, 72, 166 microtubules. Experiments performed with NuMA contained 3 nM mEGFP-dynein, 7 nM dynactin, 50 nM NuMA, 650 nM Lis1; concentrations for experiments with BicD2N$^{1\text{–}400}$ as in A. Velocities measured at 18 °C were ≈3-fold lower than those measured at 30 °C, consistent with the temperature effect previously reported (Hong *et al.*, 2016; Ruhnow, Kloß and Diez, 2017).

**Fig. S3 | Purified mScarlet-labelled NuMA C-terminal truncations. (A)** Coomassie Blue-stained SDS-PAGE of the purified NuMA C-terminal truncations. The expected molecular weight (kDa) of each NuMA construct is indicated; double bands are due to the mScarlet tag. **(B)** Comparison of Coomassie Blue staining and mScarlet fluorescent signal for samples described in A, unboiled (U) or boiled (B) in SDS sample buffer. When the protein is not boiled, it is mostly present in a state of lower apparent molecular weight (blue arrowhead), which corresponds to an mScarlet conformation preserving its fluorescence. On the contrary, upon boiling, the most abundant band is the one with higher apparent molecular weight (red arrowhead), which corresponds to a denatured mScarlet that does not fluoresce. A minor portion of protein is resistant to denaturation, which explains the double band pattern of all mScarlet-tagged NuMA proteins. **(C)** Analysis of the oligomeric state of mScarlet-tagged NuMA C-terminal constructs by mass photometry. Profiles indicate the calculated molecular weight, with the associated standard deviation (σ), of the most abundant species present in each sample. The symmetric parts of the histograms around 0 kDa represent the background signal of the buffer. In the case of NuMA$^{\text{C-term L}}$, the two peaks of almost equal height represent a mixed population of monomers and dimers.

**Fig. S4 | 1 µM NuMA$^{\text{C-term S1}}$ has only a mild effect on the dynamics of microtubules elongating from GMPCPP-seeds. (A)** Representative TIRF microscopy image (top) and kymograph (bottom) showing the growth behavior of microtubule ends (dim magenta) elongating from immobilized Atto647N-labelled GMPCPP-seeds (bright magenta) in the presence of Atto647N-tubulin and mScarlet-NuMA$^{\text{C-term S1}}$ (green). **(B)** Growth velocity distributions of minus and plus ends of microtubules under conditions explained in A (median ± 95% CI, three independent experiments). Each circle represents the growth velocity of one microtubule end segment; *n* = 101, 58 (left plot) and 109, 63 (right plot); p-values calculated from a Mann-Whitney test: 0.00 (left and right plots). **(C)** Representative TIRF microscopy image (left) and kymograph (right) showing the growth behavior of microtubule ends (dim magenta) elongating from immobilized Atto647N-labelled GMPCPP-seeds (bright magenta) in the presence of Atto647N-tubulin, before (above the dashed line) or after (below the dashed line) the addition of mScarlet-NuMA$^{\text{C-term S1}}$ (green).

**S5 | NuMA$^{\text{C-term L}}$ and NuMA$^{\text{C-term S2}}$ cap and stabilize dynamic microtubule minus ends. (A, C)** Representative TIRF-microscopy images (top) and kymographs (bottom) showing the growth behavior of microtubule ends (dim magenta) elongating from immobilized Atto647N-labelled GMPCPP-seeds (bright magenta) in the presence of Atto647N-tubulin, before (above the dashed line) or after (below the dashed line) the addition of mScarlet-labelled C-terminal NuMA fragments (green) at different concentrations. **(B, D)** Growth velocity distributions of minus and plus ends under conditions in A, C (median ± 95% CI, three independent experiments). Grey circles and gold or orange circles represent the growth before and after the addition of mScarlet-NuMA$^{\text{C-term L}}$ or mScarlet-NuMA$^{\text{C-term S2}}$, respectively. Each circle represents the growth velocity of one microtubule end segment. B: *n* = 72, 51, 66 (left plot) and 72, 53, 66 (right plot); p-values: 1.00, 0.00, 0.00 (left plot) and 1.00, 1.00, 0.19 (right plot). D: *n* = 72, 57, 72 (left and right plots); p-values: 1.00, 0.00, 0.00 (left plot) and 1.00, 1.00, 0.05 (right plot); p-values were calculated from a Kruskal-Wallis test.

**Fig. S6 | NuMA$^{FL}$ promotes nucleation in solution but does not impact γTURC-mediated nucleation.** Plots showing an increase of the solution-nucleated (A) and γTURC-nucleated (B) microtubule number over time, in the presence of 0–30 nM mScarlet-NuMA$^{FL}$ (mean ± SEM, three independent experiments). NuMA promotes spontaneous nucleation of microtubule in solution, in a dose-dependent manner (A), without exerting a clear effect on γTURC-mediated nucleation (B).

**Fig. S7 | Different modes of dynein/dynactin/NuMA$^{FL}$-mediated microtubule transport.** Representative time course TIRF microscopy images (A, C, E) and related kymographs (B, D) of microtubule transport experiments performed in the presence of 14 nM mEGFP-dynein (pre-bound to immobilized Atto647N-labelled GMPCPP-microtubules), 28 nM dynactin, 1000 nM Lis1, and "lollipop" microtubule minus-end bound mScarlet-NuMA$^{FL}$. **(A, B)** A lollipop gets aligned to the immobilized microtubule, apparently through some crosslinking dynein bound at a distance from the lollipop minus end (red arrowhead, strongest NuMA signal). The orientation is parallel, as the lollipop minus end is facing the immobilized microtubule minus end (recognized by the direction of dynein-driven transport, and accumulated dynein). **(C, D)** Similarly to A,B, a lollipop is aligned to the immobilized microtubule, however, in antiparallel orientation, which results in the plus end of the lollipop being slid towards the minus end of the immobilized microtubule. **(E)** A lollipop-bound NuMA (red arrowhead) in solution lands directly on the minus end of the immobilized microtubule, where dynein has accumulated, resulting in transport-independent gathering of minus ends.

#### VIDEO LEGENDS

**Video 1 | NuMA caps and stabilizes dynamic microtubule minus ends.** Microtubules elongate from surface-immobilized Atto647N-labelled GMPCPP-“seeds” (bright magenta), in the presence of 10 µM of Atto647N-labelled tubulin (dim magenta). Minus ends can be distinguished from plus ends as they display slower growth. After ≈10 min of imaging, 10 µM fluorescent tubulin is flowed again into the channel (white frames), together with 20 nM mScarlet-NuMA$^{FL}$ (green), which localizes selectively at the pre-elongated microtubule minus ends, arresting their growth and preventing catastrophe. Acquisition rate: 10 fps, display rate: 27 fps. Related to Fig. 4.

**Video 2 | NuMA binds to the minus ends of freshly severed microtubules.** Microtubules elongated from surface-immobilized Atto647N-labelled GMPCPP-“seeds” (bright magenta), in the presence of 10 µM of Atto647N-labelled tubulin (dim magenta) and 40 nM mScarlet-NuMA$^{FL}$ (green). Minus ends can be distinguished from plus as they are capped by NuMA and are not growing. Upon severing of the microtubules by laser ablation at multiple sites (asterisks), the longer microtubule fragments that remain attached to the surface allow the observation of both newly generated plus and minus ends. NuMA localizes and accumulates preferentially at the new minus ends. Acquisition rate: 5 fps, display rate: 24 fps. Related to Fig. 6 C.

**Video 3 | Full-length NuMA can mediate dynein/dynactin-driven microtubule transport.** Microtubule transport events observed in the presence of 14 nM mEGFP-dynein (pre-bound to immobilized Atto647N-labelled GMPCPP-microtubules), 28 nM dynactin, and 1000 nM Lis1. Microtubule “lollipops” (bright magenta) landed from solution are bound to the immobilized microtubule (dim magenta) through an anchoring point, corresponding to co-localizing mEGFP dynein (cyan) and mScarlet-NuMA$^{FL}$ (green). This results in the lollipops to dangle while being transported processively, at different speeds, towards the minus end of the immobilized microtubule. Acquisition rate: 3.5 fps, display rate: 35 fps. Related to Fig. 7 C and D.

### Figure S1

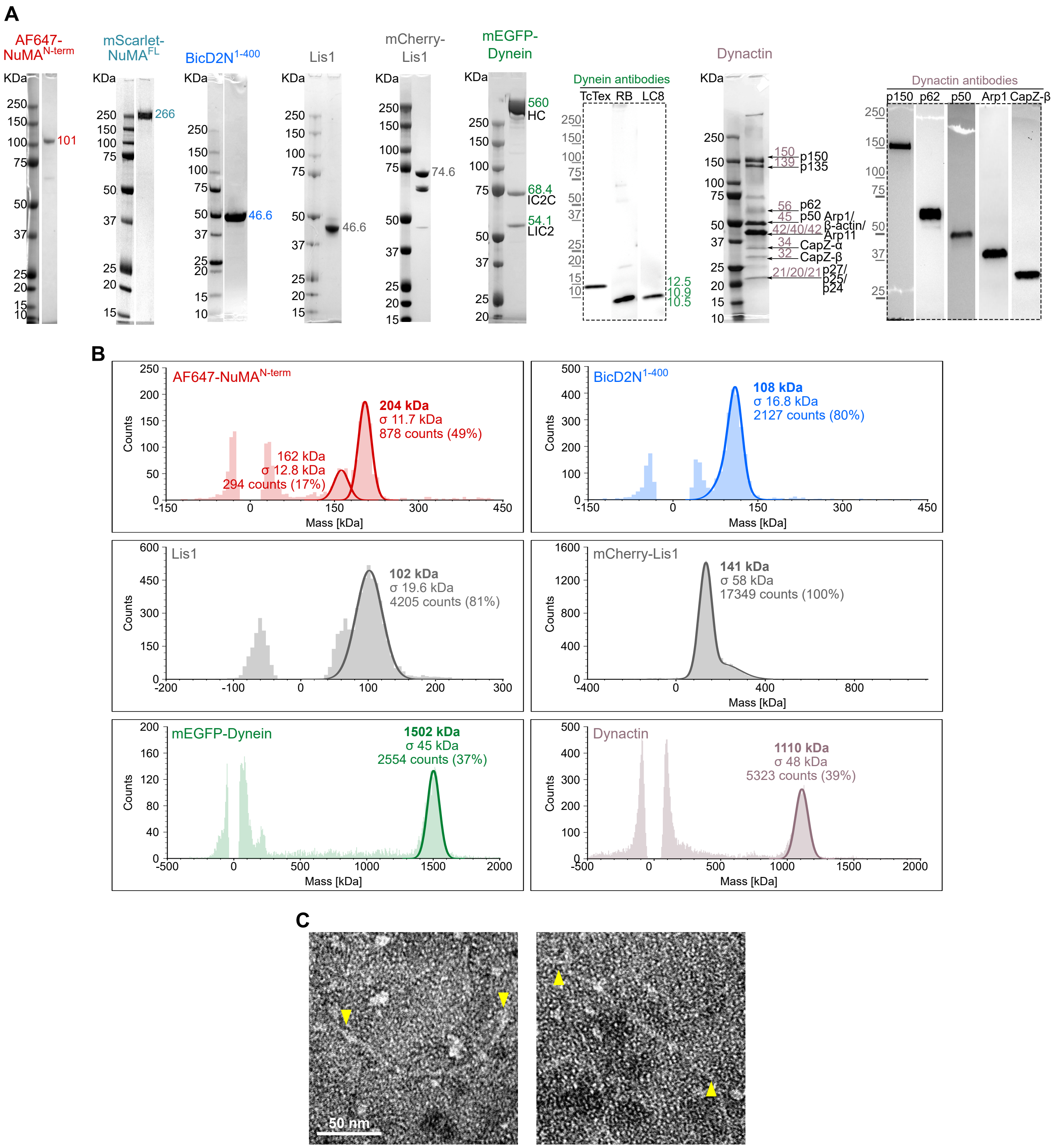

A
AF647-NuMA N-term
mScarlet-NuMA FL
BicD2N 1-400
Lis1
mCherry-Lis1
mEGFP-Dynein
Dynein antibodies
TcTex RB LC8
Dynactin
Dynactin antibodies
p150 p62 p50 Arp1 CapZ-β
KDa
101
266
46.6
46.6
74.6
560 HC
68.4 IC2C
54.1 LIC2
12.5
10.9
10.5
150 p150
139 p135
56 p62
45 p50 Arp1/
42/40/42 β-actin/ Arp11
34 CapZ-α
32 CapZ-β
21/20/21 p27/ p25/ p24
B
AF647-NuMA N-term
204 kDa
σ 11.7 kDa
878 counts (49%)
162 kDa
σ 12.8 kDa
294 counts (17%)
BicD2N 1-400
108 kDa
σ 16.8 kDa
2127 counts (80%)
Lis1
102 kDa
σ 19.6 kDa
4205 counts (81%)
mCherry-Lis1
141 kDa
σ 58 kDa
17349 counts (100%)
mEGFP-Dynein
1502 kDa
σ 45 kDa
2554 counts (37%)
Dynactin
1110 kDa
σ 48 kDa
5323 counts (39%)
Counts
Mass [kDa]
C
50 nm

### Figure S2

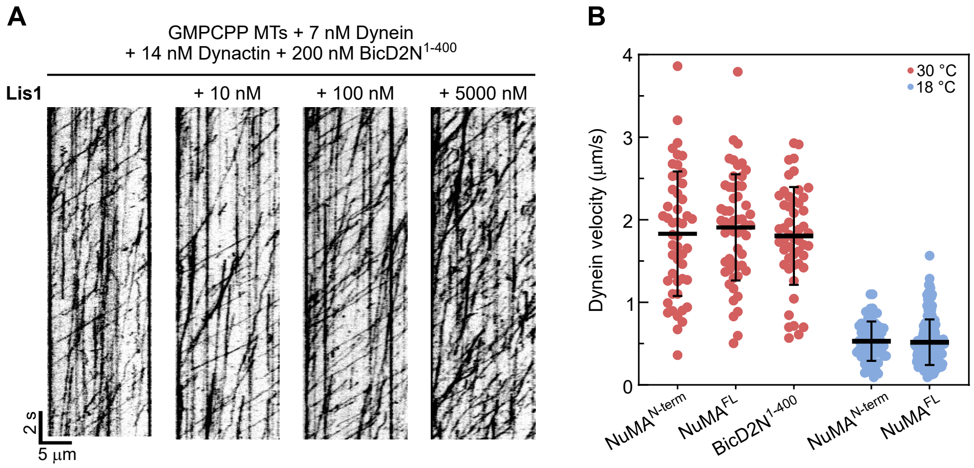

A
GMPCPP MTs + 7 nM Dynein
+ 14 nM Dynactin + 200 nM BicD2N1-400
Lis1
+ 10 nM
+ 100 nM
+ 5000 nM
2 s
5 µm
B
Dynein velocity (µm/s)
4
3
2
1
0
30 °C
18 °C
NuMAN-term
NuMAFL
BicD2N1-400
NuMAN-term
NuMAFL

### Figure S3

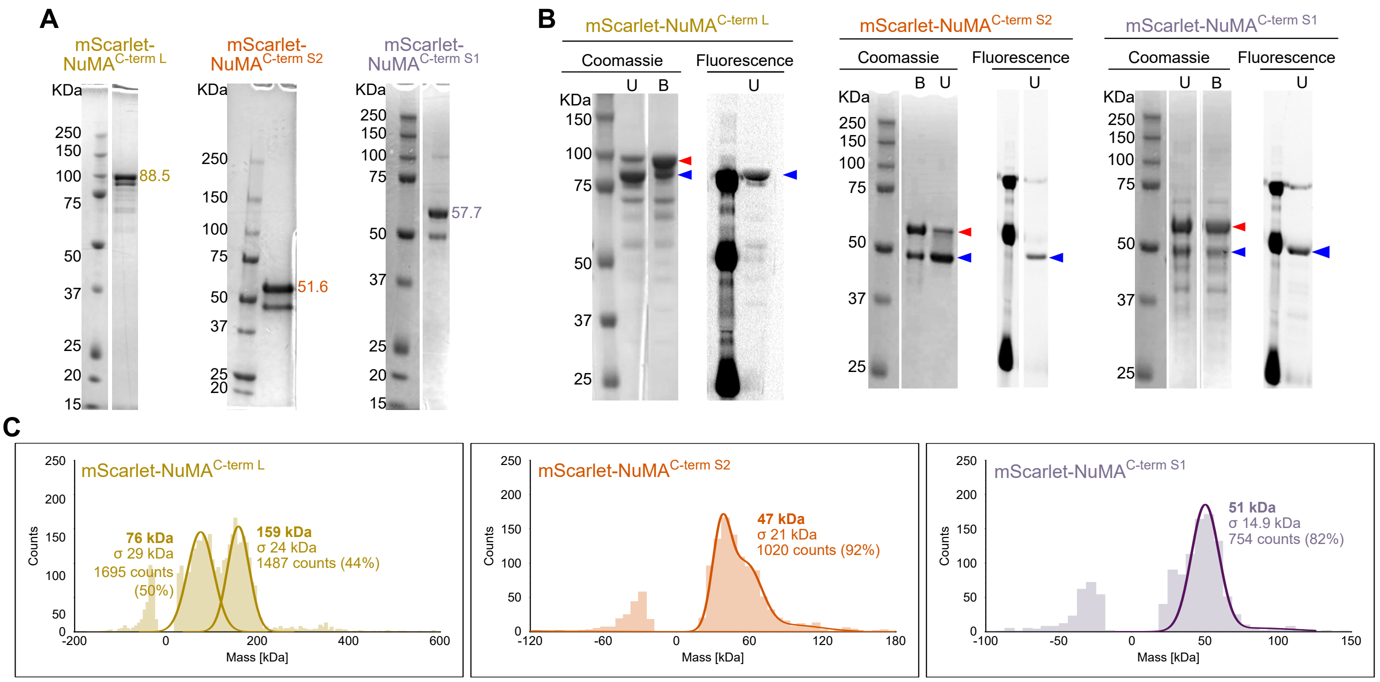

A
mScarlet-NuMA C-term L
mScarlet-NuMA C-term S2
mScarlet-NuMA C-term S1
KDa
88.5
51.6
57.7
B
Coomassie
Fluorescence
U
B
C
Counts
Mass [kDa]
76 kDa
σ 29 kDa
1695 counts (50%)
159 kDa
σ 24 kDa
1487 counts (44%)
47 kDa
σ 21 kDa
1020 counts (92%)
51 kDa
σ 14.9 kDa
754 counts (82%)

### Figure S4

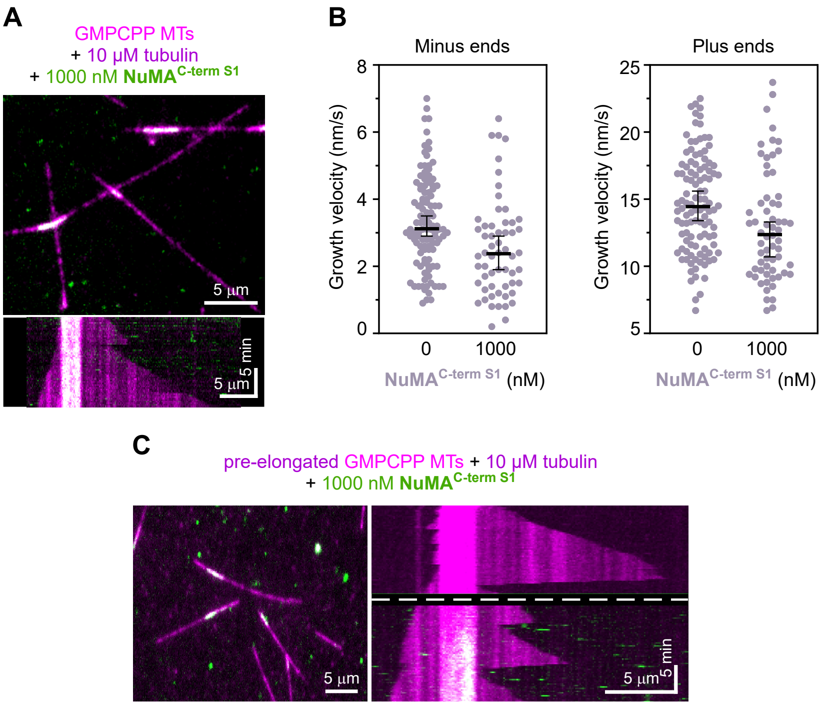

A
GMPCPP MTs
+ 10 µM tubulin
+ 1000 nM NuMA C-term S1
5 µm
5 min
5 µm
B
Minus ends
Plus ends
Growth velocity (nm/s)
0
2
4
6
8
0
1000
NuMA C-term S1 (nM)
Growth velocity (nm/s)
5
10
15
20
25
0
1000
NuMA C-term S1 (nM)
C
pre-elongated GMPCPP MTs + 10 µM tubulin
+ 1000 nM NuMA C-term S1
5 µm
5 µm
5 min

### Figure S5

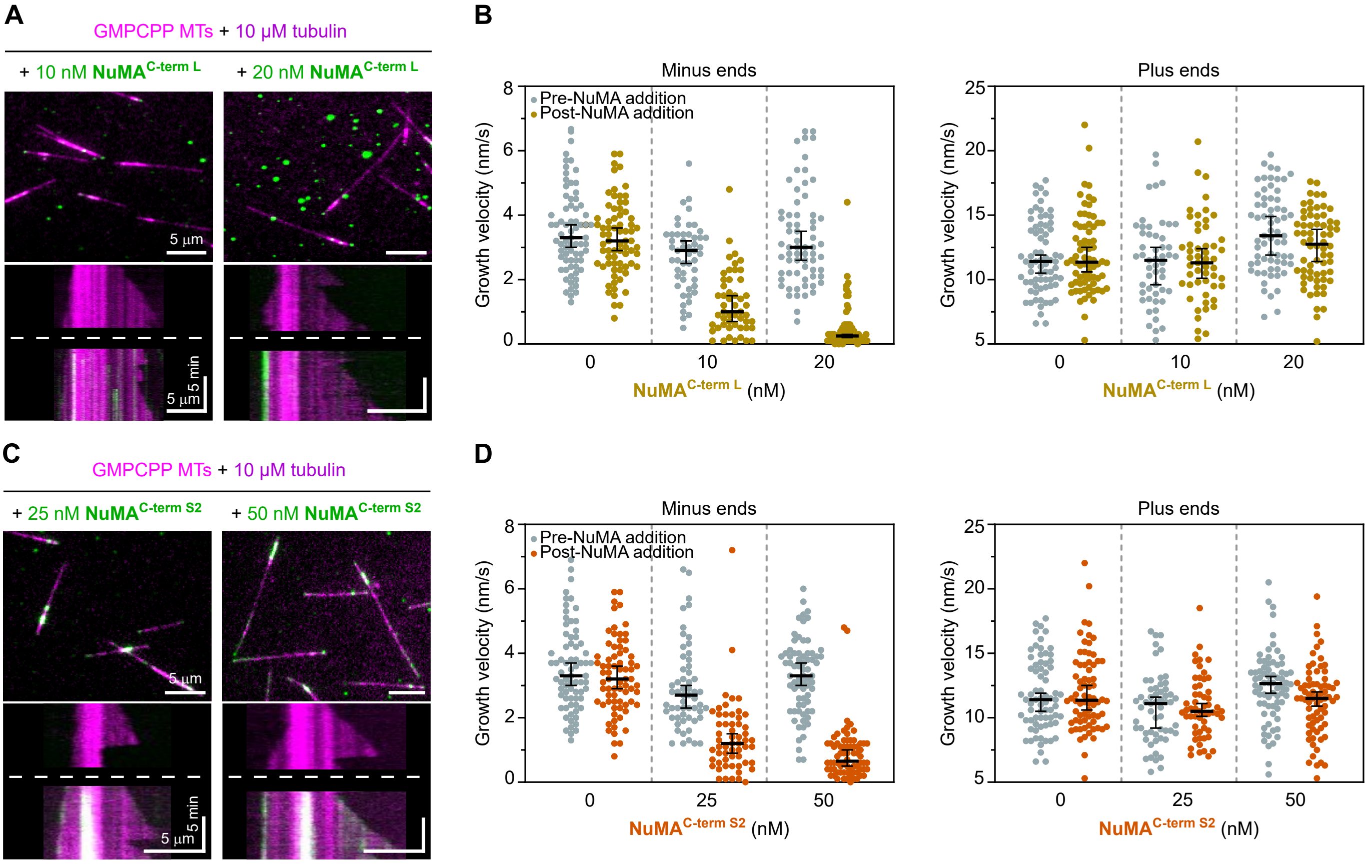
A
GMPCPP MTs + 10 μM tubulin
+ 10 nM NuMA C-term L
+ 20 nM NuMA C-term L
5 μm
5 min
B
Minus ends
Plus ends
Pre-NuMA addition
Post-NuMA addition
Growth velocity (nm/s)
NuMA C-term L (nM)
C
GMPCPP MTs + 10 μM tubulin
+ 25 nM NuMA C-term S2
+ 50 nM NuMA C-term S2
D
NuMA C-term S2 (nM)

### Figure S6

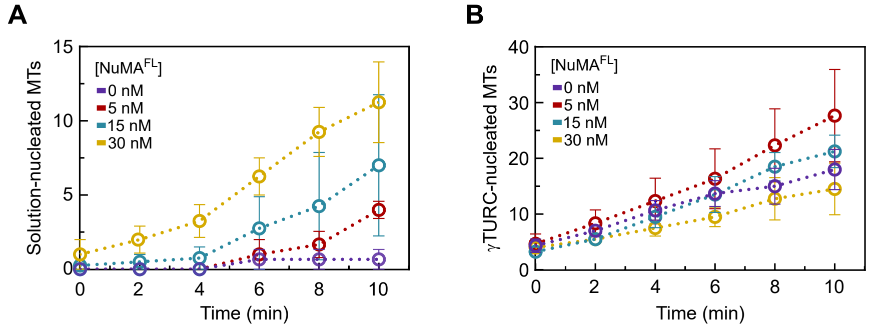
A
Solution-nucleated MTs
[NuMA^FL]
0 nM
5 nM
15 nM
30 nM
Time (min)
B
γTURC-nucleated MTs
[NuMA^FL]
0 nM
5 nM
15 nM
30 nM
Time (min)

### Figure S7

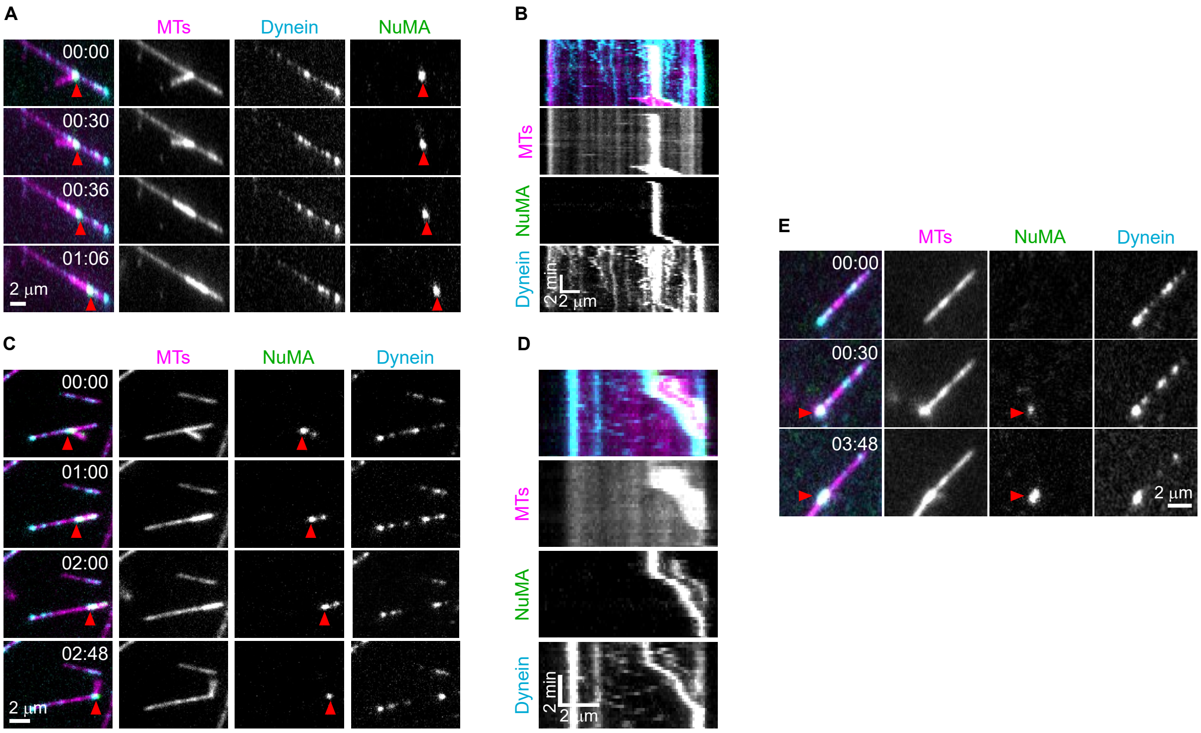
A
MTs
Dynein
NuMA
00:00
00:30
00:36
01:06
2 µm
B
MTs
NuMA
Dynein
2 min
2 µm
C
MTs
NuMA
Dynein
00:00
01:00
02:00
02:48
2 µm
D
MTs
NuMA
Dynein
2 min
2 µm
E
MTs
NuMA
Dynein
00:00
00:30
03:48
2 µm
